# Predicting Gene Spatial Expression and Cancer Prognosis: An Integrated Graph and Image Deep Learning Approach Based on HE Slides

**DOI:** 10.1101/2023.07.20.549824

**Authors:** Ruitian Gao, Xin Yuan, Yanran Ma, Ting Wei, Luke Johnston, Yanfei Shao, Wenwen Lv, Tengteng Zhu, Yue Zhang, Junke Zheng, Guoqiang Chen, Jing Sun, Yu Guang Wang, Zhangsheng Yu

## Abstract

Interpreting the tumor microenvironment (TME) heterogeneity within solid tumors presents a cornerstone for precise disease diagnosis and prognosis. However, while spatial transcriptomics offers a wealth of data, ranging from gene expression and spatial location to corresponding Hematoxylin and Eosin (HE) images, to explore the TME of various cancers, its high cost and demanding infrastructural needs significantly limit its clinical application, highlighting the need for more accessible alternatives. To bridge this gap, we introduce the Integrated Graph and Image Deep Learning (IGI-DL) model. This innovation, a fusion of Convolutional Neural Networks and Graph Neural Networks, is designed to predict gene spatial expression using HE images. The IGI-DL model outperforms its predecessors in analyzing colorectal cancer (CRC), breast cancer, and cutaneous squamous cell carcinoma (cSCC) by leveraging both pixel intensity and structural features in images. Significantly, across all cancer types, the IGI-DL model enhances the mean correlation of the top five genes by an average of 0.125 in internal and external test sets, rising from 0.306 to 0.431, surpassing existing state-of-the-art (SOTA) models. We further present a novel risk score derived from a super-patch graph, where gene expression predicted by IGI-DL serves as node features. Demonstrating superior prognostic accuracy, this risk score, with a C-index of 0.713 and 0.741 for CRC and breast cancer, supersedes traditional HE-based risk scores. In summary, the approach augments our understanding of the TME from the aspect of histological images, portending a transformation in cancer prognostics and treatment planning and ushering in a new era of personalized and precision oncology.

## Introduction

The tumor microenvironment (TME) is crucial in solid tumors’ initiation, evolution, and metastasis. Defined as a multifaceted, multicellular ecosystem, the TME conveys complex relationships among tumor cells, fibroblasts, endothelial cells, immune cells, and various extracellular matrix components^1^. An expanding amount of research has uncovered the correlation between TME and cancer patient prognosis, as well as treatment options^2,3^. Recent advancements have seen the advent of spatially resolved transcriptomics measurements aimed at revealing RNA activities in spatial terms^4^, which can characterize TME from the perspective of spatial gene expression. These methodologies employ in situ hybridization^5,6^, microdissection^7,8^, in situ sequencing^9,10^ and in situ capturing^11–13^. The commercial success of spatial transcriptomics (ST)^11^ technology by 10X Visium^14^ has significantly enhanced the application of spatially resolved transcriptomics based on in situ capturing. This technology concurrently delivers gene expression data, spatial location, and corresponding high-resolution Hematoxylin and Eosin (HE) stained histological images. Previous studies deployed 10X Visium technology to elucidate the TME across various solid tumors, investigating its contribution to tumor progression and metastasis.^15–18^.

Hematoxylin and Eosin (HE)-stained histological images, a critical component of spatial transcriptomics data, offer a wealth of information regarding tumor morphology. Thanks to the rapid advancement of deep learning (DL), numerous studies have effectively utilized these histological images to predict essential molecular biomarkers. For instance, Naik et al.^19^ employed a deep learning model to anticipate the estrogen receptor status (ERS), an important molecular biomarker for breast cancer prognosis, from HE-stained images. Schmauch et al.^20^ leveraged bulk RNA-Seq results to guide their model training, predicting gene expression of tumors from histological images. Recently, Saldanha et al.^21^ utilized swarm learning to forecast *BRAF* mutation status and microsatellite instability (MSI) from HE-stained slides of colorectal cancer. These studies verify that molecular-level information can be derived through image-level details, confirming the need for further research. In recent years, numerous studies have aimed to predict tumor patient survival using HE-stained whole slide images (WSIs) directly^22–25^. However, these survival prediction studies often fail to fully characterize the complexity of the TME at a molecular level, resulting in poor biological interpretability.

There is a clear need for a deeper and more comprehensive prognostic analysis of cancer patients based on TME characterization through spatial transcriptomics, which we seek to do. However, the cost of spatial transcriptomics has thwarted its application for survival prediction in large-scale cancer patient cohorts. HE-stained histological images, being cost-effective and easily accessible in clinical settings, present a better substitute for TME analysis. Our study, leveraging these widely available images, seeks to circumvent the current constraints of spatial transcriptomics, which is high costs and limited clinical adoption. Thus, we devised a DL model capable of predicting high-dimensional gene expression in the corresponding regions using histological images. This novel strategy allows a deeper understanding of the TME and its bearing on patient prognosis, negating the need for expensive, specialized sequencing techniques.

Some recent studies have employed DL methods to predict gene spatial expression from HE-stained images and are, therefore, the current state-of-the-art (SOTA) models^26–28^. These studies primarily exploit pixel intensity features of histological images using image-based DL techniques, such as convolutional neural networks (CNNs) and vision transformers (ViT)^29^. Gene spatial expression is intrinsically tied to the geometric information within HE-stained slides, encompassing tissue structure and cell distribution. However, these structural features cannot be adequately captured by image-based deep learning methods, leaving room for further improvement in the prediction accuracy, stability, and scalability of current methods. Graph neural networks (GNNs), adept at capturing cell structure geometries, have found extensive applications in biomedical research^30^. Cell-Graphs constructed from multiplexed immunohistochemistry (mIHC) images have demonstrated significant performance for precise survival prediction in gastric cancer^31^. A novel cell-graph convolutional neural network (CGC-Net) was proposed for grading colorectal cancer based on histology images^32^. We postulate that incorporating prior biological knowledge in nuclei segmentation and graph construction can enrich the prediction of gene spatial expression levels.

In an effort to leverage the advantages of both pixel intensity and structural features, we have designed an Integrated Graph and Image Deep Learning (IGI-DL) model. This model employs CNNs and GNNs to project HE histological slides into the gene expression space. We have tested the capabilities of our IGI-DL model across three types of solid tumors: colorectal cancer (CRC), breast cancer, and cutaneous squamous cell carcinoma (cSCC). Notably, existing SOTA models have yet to be applied to CRC, marking this as the first instance of using our unique CRC dataset to predict gene spatial expression based on histological images for this type of cancer. Furthermore, we conduct survival prediction using a constructed super-patch graph with the gene spatial expression predicted by IGI-DL as node features in the The Cancer Genome Atlas (TCGA) dataset. The IGI-DL model offers a unique perspective on spatial expression patterns and TME of solid tumors, as profiled by spatial transcriptomics, through the lens of medical image analysis. Consequently, this provides the possibility for a more accurate prognosis for cancer patients.

## Results

### Overview of IGI-DL

Figure 1 illustrates the data preprocessing workflow and the architecture for our proposed IGI-DL model. As shown in Figure 1a, the spatial transcriptome sequencing technology is used to capture the RNA from tumor samples to create the high-resolution RNA-seq gene expression map in HE-stained histological slides. There are thousands of uniquely-barcoded spots in a slide, which can capture and tag mRNAs. Each HE-stained histological image is firstly segmented into multiple non-overlapping patches according to the coordinates of each spot. We then conduct color normalization to eliminate any possible biases caused by different dyes. For each patch, we build a Nuclei-Graph where each segmented nuclei is represented as a node, and the distance of each nuclei-nuclei pair determines whether there is an edge between them. Details can be found in the section “Nuclei-Graph construction” of Methods.

**Figure 1.**
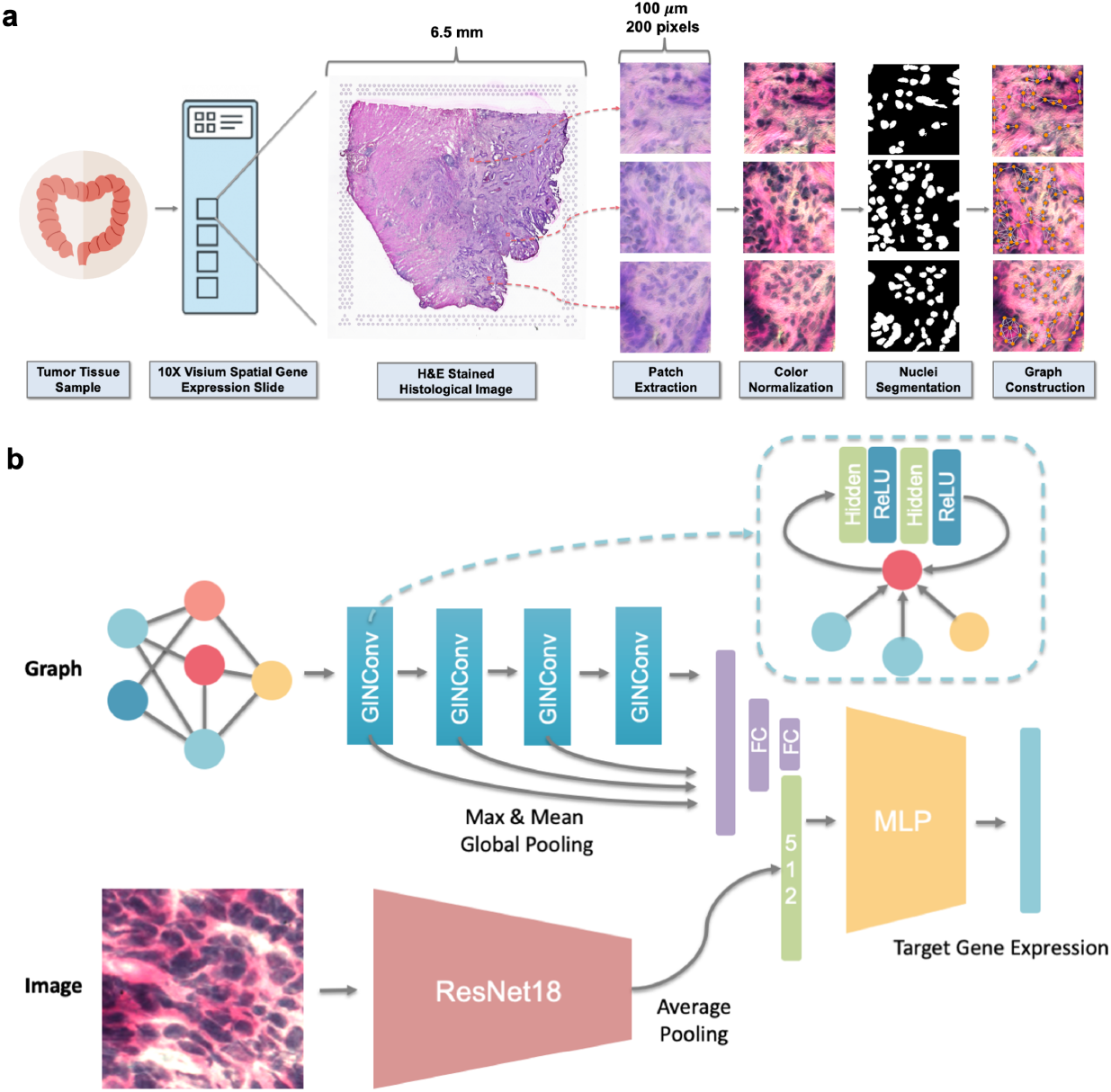
Schematic illustration of the data preprocessing workflow and our integrated graph and image deep learning (IGI-DL) architecture. **(a)** Data preprocessing workflow for model training, from HE-stained histological image to extracted color-normalized patches and constructed Nuclei-Graph. **(b)** The architecture of our designed IGI-DL model. The model has two input branches and one fusion output branch, which extract the features of the constructed Nuclei-Graph and the HE-stained image patches, and predict corresponding spatial gene expression separately.

Following the architecture shown in Figure 1b, we use IGI-DL to predict the target gene spatial expression at each spot in HE-stained images. IGI-DL takes a 200×200-pixel patch of the HE-stained histological image centered on each spot (physical size 100×100 *μm*) and the corresponding Nuclei-Graph as the input and then predicts the log-level expression of target genes, where the selection of target genes is detailed in the section “Target genes selection” of Methods.

### Performance of IGI-DL for gene expression prediction in solid tumor samples

We assess the performance of IGI-DL on the tissue samples from three different solid tumor types, including CRC, breast cancer, and cSCC. We then compare IGI-DL with three previous SOTA models, namely ST-Net^26^, HisToGene^27^, and Hist2ST^28^, which have achieved state-of-art performance in spatial gene prediction based on HE-stained histological images. We evaluate all models in the leave-one-patient-out validation set for each cancer type to ensure that the reported result can be generalized across patient samples. To further verify the generalization performance, we train the IGI-DL and other SOTA models on all patients in the leave-one-patient-out validation set and then apply the trained models to predict the gene spatial expression of tumor samples in the external test set. In the inference phase, we compute the Pearson’s correlation between the predicted and ground-truth log expression of each target gene in all spots of the tested tissue sample.

#### Colorectal cancer

For CRC, we iteratively train IGI-DL on five patients, make inferences on the remaining held-out patient in our in-house CRC dataset(Supplementary Table S1), then compare the Pearson correlation of 323 genes predicted by IGI-DL with three SOTA models. IGI-DL achieves a mean Pearson correlation of 0.161 across six held-out patients, which is significantly better than other models, with an average increase of 0.094.(Figure 2a). Comparing the model results in each patient, in most cases, our model maintains an advantage over previous models (Supplementary Figure S1). Figure 2b provides all held-out patients samples to visualize the experimental spatial expression level, and our IGI-DL predicted level for *EPCAM*, which ranks 1^*st*^ among all target genes regarding prediction results. In particular, *EPCAM* is associated with the development and metastasis of CRC and helps predict tumor stage^33^. For the CRC external test set (Supplementary Table S2), the mean correlation between the ground-truth and the IGI-DL predicted value of all target genes is 0.146, which is improved by an average of 0.101 compared to the other three models(Figure 2c). The Pearson correlation of *EPCAM* in the external sample can achieve 0.462 (Figure 2d).

**Figure 2.**
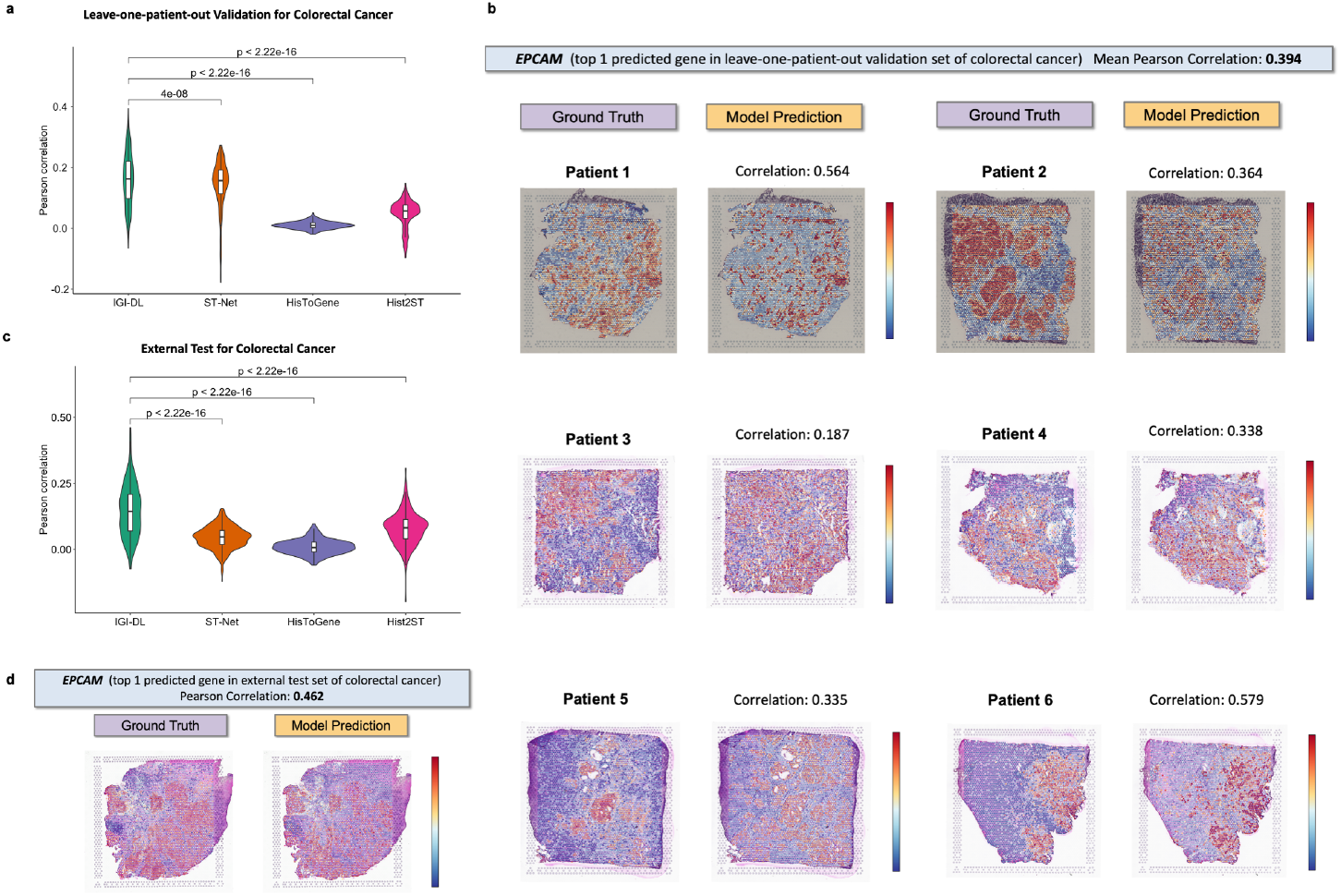
The prediction performance and visualization of gene spatial expression by our designed models in colorectal cancer (CRC). **(a)** Performance of our IGI-DL model and previous models, including ST-Net, HisToGene, and Hist2ST, for prediction of target gene spatial expression in our in-house CRC dataset. Leave-one-patient-out validation is used to calculate average performance. The evaluation matrix is mean Pearson correlation between ground-truth and predicted value of 323 target genes. The p-value is calculated by the one-tailed Wilcoxon signed rank test. **(b)** Visualization of ground-truth and predicted expression level of *EPCAM* in all tissue samples from six different patients, where *EPCAM* ranks 1^*st*^ among the prediction results of all target genes in CRC leave-one-patient-out validation set. **(c)** Comparison of our IGI-DL model with previous models for predicting gene expression in CRC external test set. **(d)** Visualization of ground-truth and predicted expression level of *EPCAM* in the tissue sample from the external test set.

#### Breast cancer

For breast cancer, we iteratively train IGI-DL on all samples from seven patients, make inferences on samples from the remaining held-out patient in the breast cancer dataset (Supplementary Table S3), and then compare the Pearson correlation of 321 genes predicted by IGI-DL with previous models. IGI-DL achieves a mean correlation of 0.200 across eight held-out patients. The violin plot in Figure 3a shows that our IGI-DL model outperforms all SOTA models, with an average increase of 0.063. When comparing these models at the level of each patient and each sample separately, our model performs best in most cases (Supplementary Figure S2, and S3). *ERBB2*, an oncogene in breast cancer^34^, ranks 1^*st*^ among all target genes in breast cancer leave-one-patient-out validation set with a mean correlation of 0.374 (Supplementary Figure S4, S5, S6, and S7).

**Figure 3.**
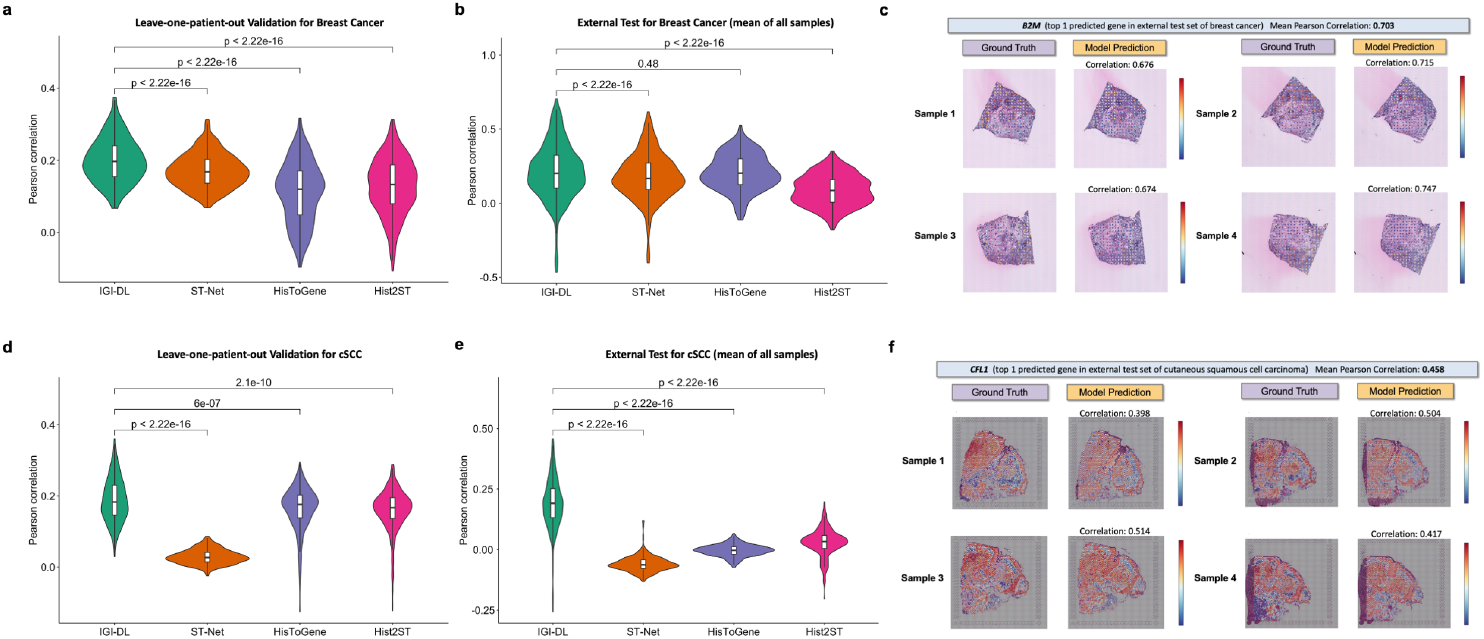
Comparison of gene expression prediction performance among different models and visualization of gene spatial expression by our designed models in breast cancer and cutaneous squamous cell carcinoma (cSCC). **(a)** Violin plot comparing the mean Pearson correlation of target genes prediction among our model and previous SOTA models in breast cancer leave-one-patient-out validation set. The p-value is calculated by the one-tailed Wilcoxon signed rank test. **(b)** Violin plot comparing the mean Pearson correlation of target genes prediction among our model and previous models in the breast cancer external test set. **(c)** Visualization of ground-truth and predicted expression level of *B2M* by IGI-DL in four tissue samples from breast cancer leave-one-patient-out validation set, where *B2M* ranks 1^*st*^ among the prediction results of all target genes. **(d)** Violin plot comparing the mean Pearson correlation of target genes prediction among our model and previous models in the cSCC leave-one-patient-out validation set. **(e)** Violin plot comparing the mean Pearson correlation of target genes prediction among our model and previous models in the cSCC external test set. **(f)** Visualization of ground-truth and predicted expression level of *CFL1* by IGI-DL in four tissue samples from cSCC leave-one-patient-out validation set, where *CFL1* ranks 1^*st*^ among the prediction results of all target genes.

Of the 321 predicted genes in the leave-one-patient-out validation set, 317 genes exist in the breast cancer external test set. The Pearson correlation of these genes predicted by IGI-DL is 0.213, significantly higher than that of ST-Net and Hist2ST at the patient and sample level, while HisToGene performs similarly to our IGI-DL model (Figure 3b, and Supplementary Figure S8). Figure 3c visualizes the experimental spatial expression level, and the predicted level for the top 1 gene *B2M*, which achieves a mean correlation of 0.703 for external test samples. In particular, *B2M* plays a multifaceted role in cancer immunotherapies^35^.

#### Cutaneous squamous cell carcinoma

For cSCC, we iteratively train IGI-DL on all samples from three patients and make inferences on samples from the remaining held-out one patient in the cSCC dataset (Supplementary Table S4). We compare the Pearson correlation of 487 genes predicted by IGI-DL with previous models. IGI-DL achieved a mean correlation of 0.188 across four held-out patients, which has the best performance among all models, with an increase of 0.129 more than the average performance of other SOTA models (Figure 3d). Comparing the level of each patient and each sample separately, our IGI-DL model still performs best in most cases (Supplementary Figure S9). *KRT5*, one of the cancerous squamous cell-associated genes^36^, ranks 1^*st*^ among all target genes in cSCC leave-one-patient-out validation set with a mean correlation of 0.361 (Supplementary Figure S10, and S11).

Of the 487 predicted genes in the leave-one-patient-out validation set, 467 genes also exist in the cSCC external test set. For four samples in the cSCC external test set, the mean correlation between the ground-truth and the IGI-DL predicted value of all target genes is 0.191, which is significantly higher than that of ST-Net, HisToGene and Hist2ST at the patient and sample level. (Figure 3e, and Supplementary Figure S12). Figure 3f visualizes the ground-truth and predicted expression pattern for top 1 gene *CFL1* in the external test set, achieving a mean correlation of 0.458. Studies have already shown that *CFL1* can control actin stress fibers, nuclear integrity, and cell survival, which is related to the development of cSCC^37^.

### IGI-DL outperforms previous SOTA models

Table 1 lists the top five genes which can be best-predicted by IGI-DL, with a much higher Pearson correlation than that predicted by previous SOTA models. For the validation set of CRC, the top five genes mean correlation of our IGI-DL can achieve 0.374, with an improvement of 0.113 compared to the best SOTA model (ST-Net) under this condition. In terms of the external set of CRC, our IGI-DL attains the top five genes mean correlation of 0.424, which outperforms the best SOTA model (Hist2ST) in this case by an improvement of 0.172. In the validation set for breast cancer, our IGI-DL demonstrates the top five genes mean correlation of 0.367, improving by 0.066 compared to the best SOTA model (ST-Net) in this scenario. In the external test set for breast cancer, the top five genes mean correlation of our IGI-DL is 0.641, indicating a 0.076 improvement over the best SOTA model (ST-Net). When tested on the validation set for cSCC, our IGI-DL performs with the top five genes mean correlation of 0.343, surpassing the best SOTA model (His2ST) by 0.061. When further tested on the external test set for cSCC, our IGI-DL achieves the top five genes mean correlation of 0.436, representing a 0.259 improvement over the best SOTA model (Hist2ST). The best SOTA model is not fixed in the internal validation and external testing sets of different tumors, but our IGI-DL model’s performance are always better than the best result of other models, with an average improvement of 0.125 for the top five genes mean correlation. Our IGI-DL model has shown excellent performance in various types of solid tumor, surpassing other models (Supplementary Table S5). Moreover, its performance on the external test samples demonstrates good generalization ability without overfitting.

**Table 1.**
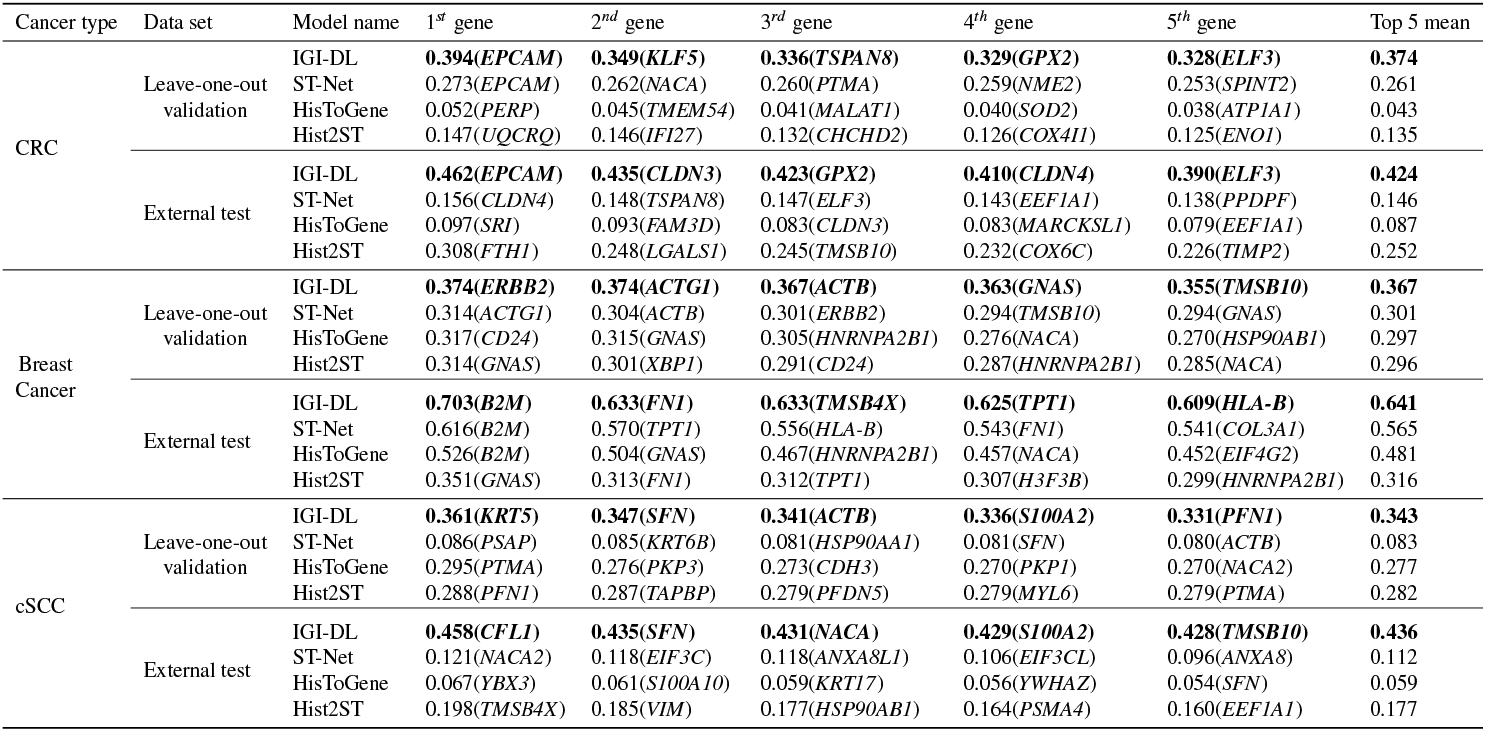
Pearson correlation between the predicted and ground-truth gene expression for the top five predicted genes of our proposed IGI-DL and previous models

### Pathway and process enrichment analysis based on best-predicted genes by IGI-DL

Supplementary Figure S13 presents pathway and process enrichment analysis results based on genes most accurately predicted by IGI-DL in CRC. The top-ranking pathway is “Oxidative phosphorylation”, a prognostic marker for CRC^38,39^. “Cell-cell adhesion”, a key factor in cancer development and metastasis^40^, is the second most significant process. Ranking third is “Positive regulation of cell motility”, essential for metastatic tumor cell migration^41^. The fourth-ranking process is “Epithelial cell differentiation”; Notably, the loss of this differentiation is seen at tumor invasion fronts and influences metastasis^42^.The fifth pathway is “Estrogen-dependent gene expression” is the top 5 pathway. Global data suggests a lower CRC incidence in women than men, indicating estrogen’s protective role against CRC^43^. All these pathways and processes, enriched based on significant p-values, are intrinsically linked to progression, metastasis, and prognosis of CRC.

Regarding breast cancer, Supplementary Figure S14 displays enrichment analysis results for pathways and processes using the most accurately predicted genes. The top process is “Antigen processing and presentation of endogenous antigen”, associated with immune evasion in breast cancer and potential therapeutic targeting^44^. The second-ranking pathway is the “VEGFA-VEGFR2 signaling pathway”, a potential target for breast cancer treatment that could inhibit cancer angiogenesis when suppressed^45,46^. Ranking third is the “Neutrophil degranulation” pathway, with recent research demonstrating the role of tumor-associated neutrophils in immunosuppressive process of the breast cancer’s TME^47^. The fourth-ranked process is “Axon guidance”, where axon guidance molecules are tumor suppressors and oncogenes in breast cancer^48^. The fifth-ranking pathway is “Scavenging by Class A Receptors”. Of note, Scavenger receptor class A member 5 (SCARA5), involved in this pathway, plays a pivotal role in breast cancer progression and metastasis^49^.

Supplementary Figure S15) illustrates the pathway and process enrichment analysis based on genes most accurately predicted by IGI-DL for cSCC. The top-ranking pathway “Neutrophil degranulation”. Notably, molecular pathway aberrations, including neutrophil degranulation signaling pathways, have been identified as potential drug targets in invasive and metastatic cSCC^50^. The second most significant pathway is “RHO GTPase Effectors”. Specifically, Rho GTPase signaling networks are known to contribute to skin cancer progression^51^. Ranking third among all enriched pathways and processes is “Formation of the cornified envelope”, which indicates cell death in the skin^52^. Intriguingly, a recent study that conducted a Reactome analysis of all differential proteins highlighted common pathways involved in the development of cSCC^53^, with “Neutrophil degranulation” and “Formation of the cornified envelope” being significantly represented and also top-ranking in our study. The fourth top-ranking pathway is “VEGFA-VEGFR2 signaling pathway”. Therapies targeting VEGFA/VEGFR2 can thwart VEGFA-induced regulatory T-cell proliferation in cancer^54^. “Cell-cell adhesion” ranks fifth among all enriched pathways and processes. Cell Adhesion Molecule 1 (CADM1), which participates in this process, is an independent prognostic factor for patients with cSCC^55^.

The enrichment analysis of pathways and processes based on genes most accurately predicted in various cancers demonstrate that the expression levels of genes tied to the TME, tumor cell growth, and metastasis can be effectively inferred from tissue morphology and cellular structure in histological images.

### Ablation experiment for IGI-DL

To explore the role of integrating geometric and texture features in the architecture design for IGI-DL, we conduct the ablation experiment by comparing the performance of image and graph based models with integrated models. As demonstrated in Figure 4a and Supplementary Table S6, we use leave-one-out cross-validation in our in-house CRC dataset to evaluate the performance of different models. For predicting 323 target genes, IGI-DL has the best performance, with a mean correlation of 0.161 across all held-out patients. In 202 of 323 target genes, the expression values predicted by IGI-DL correlate positively with experimental ground-truth values at the 0.05 significance level for all six held-out patients. Furthermore, IGI-DL has 54.49% well-predicted genes among all target genes with a mean correlation across all held-out patients larger than 0.15. Considering these three evaluation matrics comprehensively, we can conclude that integrated models have a higher overall performance than image and graph based models.

**Figure 4.**
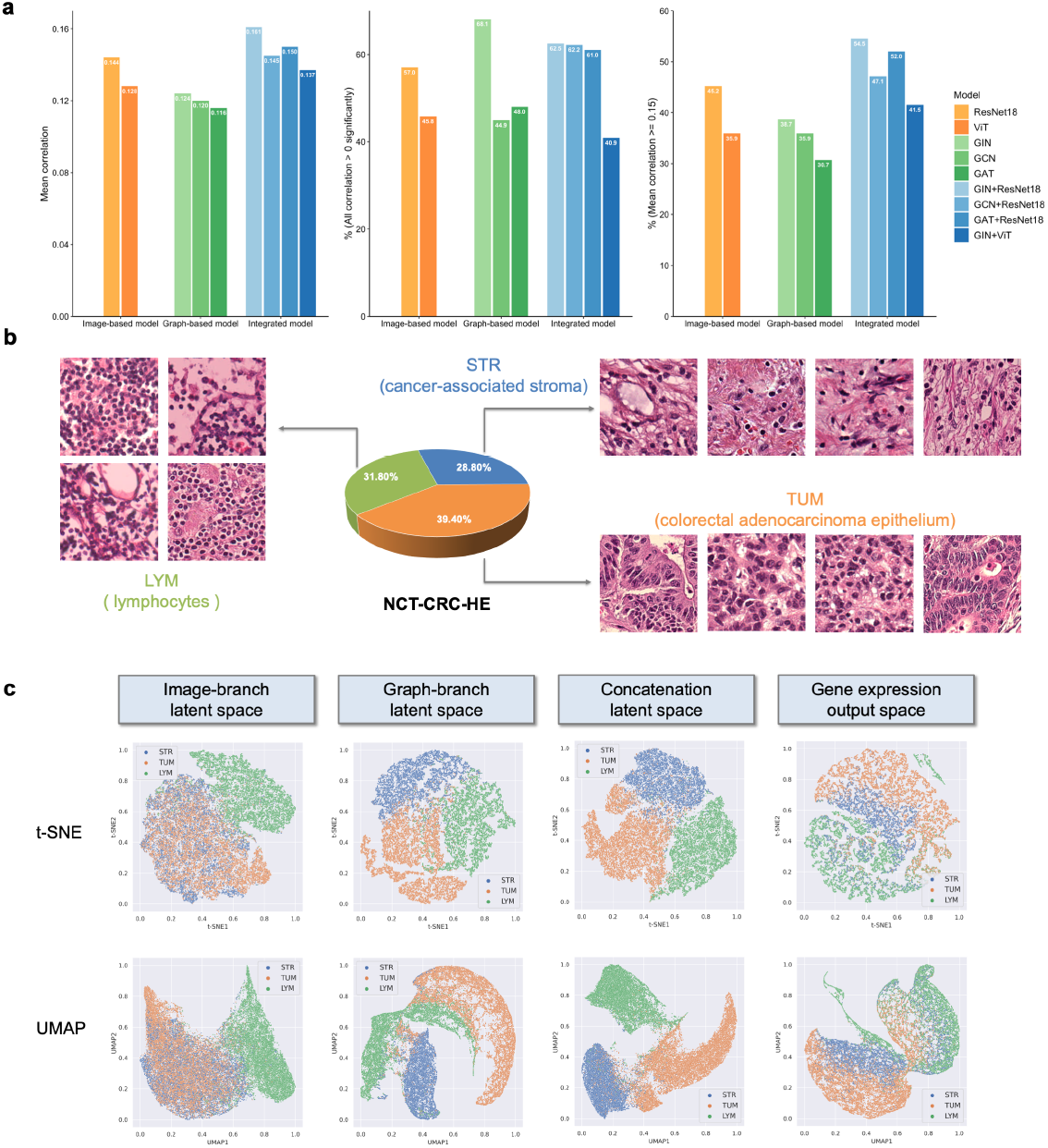
The ablation experiment and 2-D visualization of different latent spaces in our IGI-DL model for colorectal cancer. **(a)** Performance of image-based, graph-based and integrated models for prediction of target gene spatial expression in our in-house CRC dataset. Left evaluation matrix is mean Pearson correlation between ground-truth and predicted value of 323 target genes. Middle evaluation matrix is percentage of genes whose predicted values correlate positively with experimental ground-truth value at 0.05 level. Right evaluation matrix is percentage of genes with correlation larger than 0.15. **(b)** Typical patches with different labels in NCT-CRC-HE dataset. LYM represents patches with many lymphocytes, STR represents patches of cancer-associated stroma and TUM represents patches of colorectal adenocarcinoma epithelium. **(c)** t-SNE and UMAP dimensionality reduction visualization of feature vectors in different parts of the IGI-DL, from two input branches to the output branch. Green, blue and orange points indicate LYM, STR and TUM patches, respectively.

Among different integrated models, IGI-DL, with four layers of GIN as the graph features extractor and ResNet18 as the image features extractor exhibits the best performance. Supplementary Table S7 lists the top five genes predicted by IGI-DL and then compares the performance with other models. These results of the ablation experiment suggest that the combination of texture features and geometric information in the tissue morphology extracted by CNN-based image-branch and GNN-based graph-branch in IGI-DL can boost the performance for predicting gene spatial expression.

### Latent space visualization of IGI-DL

To explore the extracted features of different hidden layers and output gene expression in IGI-DL, we use t-distributed stochastic neighbor embedding (t-SNE)^56^ and uniform manifold approximation and projection (UMAP)^57^ to project latent space into low-dimensional subspaces. IGI-DL with weights trained on all patients in our in-house CRC dataset is applied to the NCT-CRC-HE dataset. As shown in Figure 4b, the public dataset has typical patches in CRC HE-stained images with different labels, including lymphocytes, cancer-associated stroma, and colorectal adenocarcinoma epithelium.

Figure 4c demonstrates latent space visualization of feature vectors in different parts of IGI-DL, from two input branches to the output branch. Visualization using t-SNE and UMAP shows a similar pattern. Image-branch latent space represents the output of ResNet18 in IGI-DL. In this subspace, lymphocytes patches cluster separately, but stroma and tumor patches are mixed. Graph-branch latent space is the output of four layers of GIN in IGI-DL. Three types of patches form three separate clusters in this low-dimensional subspace. Concatenation latent space combines the extracted image features and graph features, where three types of patches are relatively scattered among each other. In gene expression output space, lymphocytes patches are grouped separately, and stroma patches surround tumor patches. The distribution pattern of stroma and tumor patches in latent space is consistent with their mutual location in the TME, where cancer-associated stroma is in the adjacent area around the solid tumor. Latent space visualization shows the gene expression of lymphocytes, stroma, and tumor regions predicted by IGI-DL has similar low-dimensional distribution patterns compared with the previous study’s single-cell sequencing results^58^.

Tumor-stroma interactions can determine the biological behavior of tumor growth, infiltration, and metastasis^59^. The gene spatial expression patterns predicted by our IGI-DL can help further reveal the TME from the perspective of histological images.

### Prognosis prediction with a super-patch graph

We apply IGI-DL to infer gene spatial expression from HE-stained histological images, construct a super-patch graph, and then conduct prognosis prediction in the TCGA-CRC (Supplementary Table S8) and TCGA-BRCA cohort (Supplementary Table S9), which contains both WSIs and survival data of CRC and breast cancer patients. As shown in Figure 5a, our graph-based survival model predicts the super-patch graph risk score for each patient. We compare the performance of graph-based survival models using super-patch graphs constructed by different patch feature extraction methods, including spatial gene expression predicted by IGI-DL, and features extracted by DenseNet^60^ and ResNet^61^. The box plot in Figure 5b illustrates that using spatial gene expression predicted by IGI-DL as node features in the super-patch graph can boost the performance of our survival model for both CRC and breast cancer. In the TCGA-CRC cohort, the survival model based on super-patch graphs with spatial gene expression as node features can achieve a mean C-index of 0.713 in five-fold cross-validation, which is higher than 0.649 and 0.677 for super-patch graphs with DenseNet and ResNet features. In the TCGA-BRCA cohort, the survival model based on super-patch graphs with spatial gene expression as node features can achieve a mean C-index of 0.741 in the five-fold cross-validation, higher than 0.705 and 0.679 for super-patch graphs with DenseNet and ResNet features (Supplementary Table S10).

**Figure 5.**
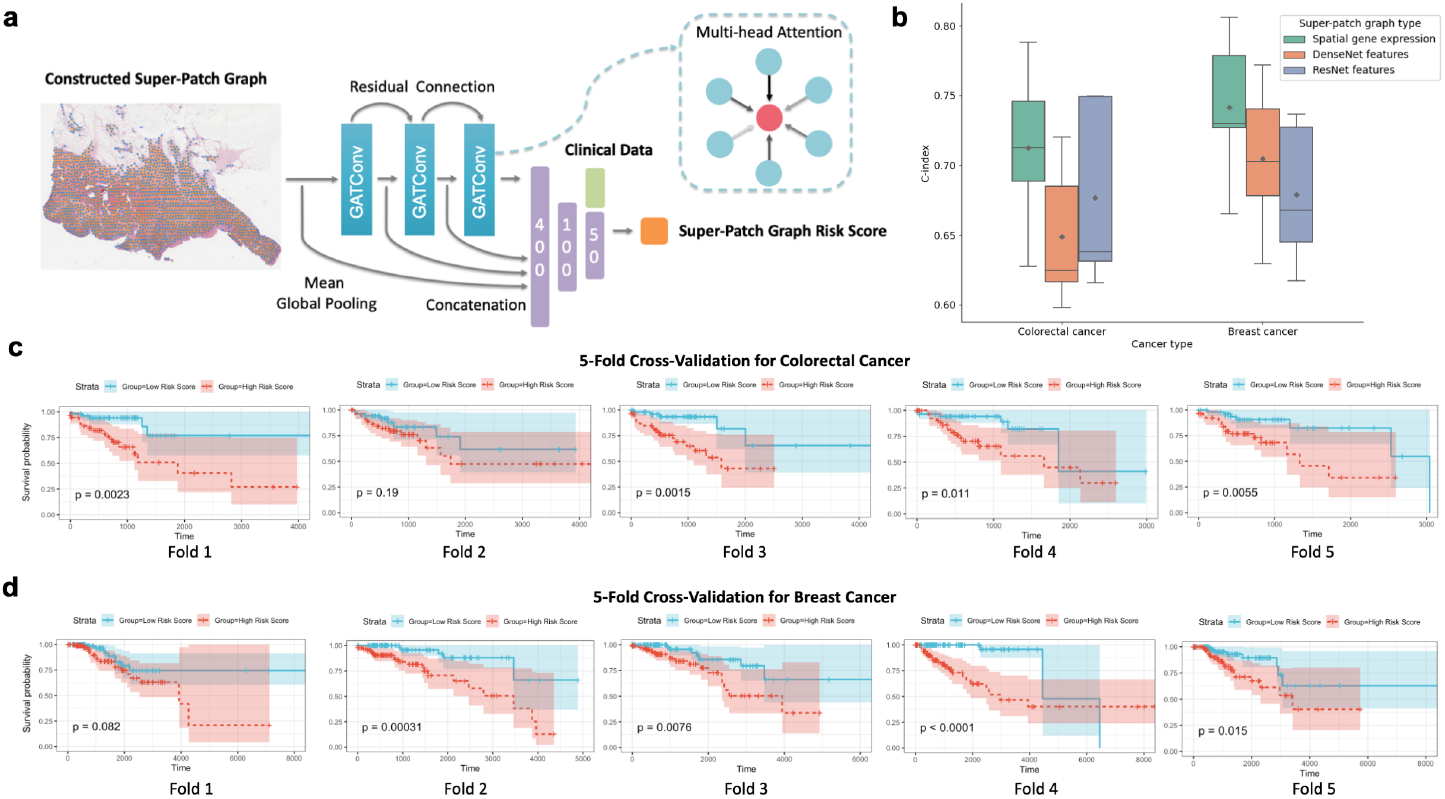
Graph-based survival prediction in TCGA-CRC and TCGA-BRCA cohorts. **(a)** The architecture of our graph-based survival model. The model takes the constructed super-patch graph as input and uses three layers of GAT to extract features from the graph. In the second-to-last layer of the model, these extracted graph features are combined with clinical data, and the final output is the patient-level risk score. **(b)** Box plot comparing the performance of graph-based survival models using super-patch graphs constructed by different patch feature extraction methods in colorectal cancer and breast cancer. The Y-axis represents C-index and the results of each method are obtained through five-fold cross-validation. Green represents using our IGI-DL model to extract patch-level gene expression features. Orange and blue represent using pre-trained DenseNet and ResNet to extract patch features, respectively. **(c)** Kaplan-Meier plots of patients’ overall survival with regard to super-patch graph risk score for each validation fold in the five-fold cross-validation of colorectal cancer, where risk score is the output of the survival model using super-patch graphs constructed by patch-level gene expression features predicted by IGI-DL. Patients are divided into high-risk group (indicated in red) and low-risk group (indicated in blue) according to the median risk score. **(d)** Kaplan-Meier plots of patients’ overall survival with regard to super-patch graph risk score for each validation fold in the 5-fold cross-validation of breast cancer, where risk score is the output of the survival model using super-patch graphs constructed by patch-level gene expression features predicted by IGI-DL.

Figure 5c and 5d show Kaplan-Meier plot of patients’ overall survival concerning risk score based on the super-patch graph with spatial gene expression features in the test set of each fold for CRC and breast cancer, respectively. In most test folds, patients in the high-risk group (super-patch graph risk score is larger than the median risk score) show significantly shorter overall survival than those in the low-risk group. These results indicate that the predicted super-patch graph risk score can be an independent prognostic indicator for CRC and breast cancer patients.

Based on the trained graph-based survival model, we can identify some genes that are either favorable or unfavorable for prognosis based on the positive or negative scores obtained through the Integrated Gradients (IG) method^62^. For all tested patients in the TCGA-CRC cohort, the IG score values for gene *KLF5* are all positive, indicating that the upregulation of *KLF5* expression in each super-patch would increase the patient’s risk score, which is consistent with previous research findings^63,64^. On the contrary, the IG score values for gene *GAS5* in CRC patients are all negative, indicating this is a prognostic protective gene. Previous studies also showed that gene *GAS5* could inhibit CRC cell proliferation^65–67^. If only the predicted spatial expression of these two genes in tissue is visually analyzed, there is no significant difference between patients with good and poor prognosis (Supplementary Figure S16), highlighting the importance of constructing a graph structure including multiple gene spatial expression levels as input for prognosis prediction models.

We obtained similar results during the analysis of the trained graph-based survival model for breast cancer using the IG method (Supplementary Figure S17). For all tested patients in the TCGA-BRCA cohort, the IG score values for gene *HNRNPA2B1* are all positive, indicating that the expression of this gene is unfavorable for survival. Related research have suggested that *HNRNPA2B1* can promotes the proliferation of breast cancer cells and contribute to the tumorigenic potential of breast cancer cells^68,69^. Conversely, the IG score values for gene *PFN1* in breast cancer patients are all negative, which means that the upregulation of *PFN1* expression in each super-patch would decrease the patient’s risk score. *PFN1* is a negative regulator of breast cancer aggressiveness, and its overexpression can inhibits proliferation of breast cancer cells^70–72^. Predicting gene expression in different tissue regions based on pathological images is beneficial for the overall characterization of the TME and promotes precise prognosis of cancer patients.

## Discussion

In the rapidly expanding healthcare data landscape, our research focuses on predicting the spatial expression level of numerous target genes using high-resolution HE-stained histological images, achieved through our integrated deep learning model, IGI-DL. The fusion strategy and specific architecture of IGI-DL directly enhance prediction performance compared to models reliant on either graphs or images alone. Through these techniques, the GIN capably captures the morphological characteristics of nuclei and their associations with peripheral cells from Nuclei-Graphs, whereas ResNet18, trained from scratch, extracts extracellular matrix features and local tissue environmental conditions from the image pixel matrix.

Significant differences exist in the number and size of spots in histological slides among ST and 10X Visium. For CRC and breast cancer, the spatial transcriptomics platform used for internal validation set and external test set is consistent, 10X Visium and ST, respectively. For cSCC, ST is used for the internal validation set while 10X Visium is used for the external test set. Under this difference between internal and external data set for cSCC, the performance of our IGI-DL is still strong, showing good generalization ability. ST-Net^26^ was initially designed for breast cancer spatial transcriptomics obtained through ST platform. Our results shows that ST-Net can achieve good performance in both internal validation and external test set of breast cancer, which is only slightly lower than IGI-DL model. For CRC, ST-Net performs well on the internal validation set, but does not generalize to the external test set. For cSCC, ST-Net has unsatisfactory performance. HisToGene^27^ and Hist2ST^28^ are similar to each other, using ViT or GNN to model the whole slice in spatial transcriptomics, which are more applicable when the number of spots per slide is small, such as in ST platform. For the cSCC internal validation set obtained by ST platform, both HisToGene and Hist2ST perform well, inferior to IGI-DL method but far better than ST-Net. However, in the cSCC external test set obtained by 10X Visium, these two models perform poorly, and the mean correlation of HisToGene is even negative. To sum up, compared with current SOTA models, IGI-DL demonstrates a superior and stable performance in multiple solid tumor types, and also has excellent cross-platform generalization ability.

The TME contains many different cell types, and the distribution of gene expression varies among different cell types. Although ST and 10X Visium has not reached the resolution of single-cell, it has been able to accurately characterize different regions such as tumor cells, immune cells, and tumor stroma from the perspective of gene expression in spots. Gene spatial expression predicted by IGI-DL based on patches with different labels in NCT-CRC-HE shows a unique distribution in the low-dimensional subspace. The distribution of gene expression inferred from tumor and immune patches is relatively separate, consistent with the fact that receptors and ligands in immune checkpoints come from tumor cells and immune cells, respectively.

The gene expression inferred from cancer-associated stroma patches is distributed in the surrounding area of the tumor patches, which is also consistent with the actual physical distribution of stroma and cancer regions in the tissue. IGI-DL can effectively capture the distribution of stromal cells and immune cells in tumor tissues and characterize TME.

Interestingly, in different solid tumor types, the most significant pathways enriched by the accurately predicted genes based on the IGI-DL are all associated with the corresponding tumors’ incidence, metastasis, and prognosis. This further confirms that the texture information and cellular topology structure in histological images can effectively reveal gene expression levels related to tumor development. In particular, for cSCC, the enrichment analysis results based on accurate predicted genes were consistent with those of enrichment analysis on differential proteins^53^. It is clinically significant that accurate prediction of these gene expression levels contributes to the precise prognosis of patients. Our IGI-DL has become an effective bridge connecting the information at the image level and molecular levels. Constructing super-patch graphs from WSIs can depict the overall topological structure and heterogeneous distribution of the tumor tissues. Existing research typically extract and vectorize features of each patch using a DL model, VGG^73^ or ResNet, pre-trained on natural images dataset such as ImageNet^22–25^, which has a certain level of representational power but lacks interpretability in the biomedical field. Using our IGI-DL, it is possible to embed molecular-level information, gene spatial expression level, in the corresponding patch region into the super-patch graph structure. The results on TCGA-CRC and TCGA-BRCA both indicate that the risk score obtained through a super-patch graph with gene expression predicted by IGI-DL as node features can achieve more accurate prognosis for tumor patients.

However, we do acknowledge the potential limitations of our study. Primarily, our model is designed for the most widely used spatial transcriptomics platforms, ST and 10X Visium, without incorporating platforms such as MERFISH and Slide-seq. Moreover, our model has yet to achieve super-resolution prediction. Looking forward, we are exploring the extension of gene spatial expression and prognosis prediction to more solid tumors. Ultimately, we aspire to leverage histological images and spatial gene expression to infer the evolutionary trajectories in tumor tissues, thereby gaining a more profound understanding of the mechanisms underlying solid tumor onset and progression.

IGI-DL is an effective integrative tool that couples CNN and GNN to fully explore tissue texture information and spatial cell structure in histological images, accurately predicting the spatial expression of critical genes. Additionally, the risk score predicted by the super-patch graph with gene expression values as node features could serve as a new prognostic biomarker, aiding in patient-specific treatment plans and ultimately facilitating precision medicine. Our study lays down a blueprint for future efforts in this domain, pushing the frontier of deep learning applications in computational pathology and, more broadly, healthcare.

## Methods

### Dataset

This study utilised spatial transcriptomics data from three different cancer types for modeling and evaluation, including CRC, breast cancer, and cSCC. For colorectal cancer, we used spatial transcriptomics data from six colorectal cancer patients in Ruijin Hospital, Shanghai Jiao Tong University sequenced by 10X Visium as a leave-one-patient-out validation set (Supplementary Table S1). We utilized a publicly available 10X visium sequencing sample from a colorectal cancer patient in the previous study^74^ as an external test set (Supplementary Table S2). For breast cancer, we used a publicly available breast cancer dataset published in previous research^75^ as our leave-one-patient-out validation set. The dataset contained 36 tissue regions from eight patients and was sequenced using ST technology. We used another public ST technology sequenced breast cancer dataset^11^ as our external test set, which contains four tissue regions from one patient (Supplementary Table S3). For cutaneous squamous cell carcinoma, we used a publicly available ST sequenced cSCC dataset published in previous research^15^ as our leave-one-patient-out validation set, including 12 tissue regions from four patients. Another public 10X Visium sequenced cSCC dataset^76^, including four tissue regions from one patient, were used as the external test set (Supplementary Table S4). The histological images of these tissue regions are all stained with Hematoxylin and Eosin (HE). For ST technology, the distance between the adjacent spots is 200 *μm*, and the diameter of each spot is 100 *μm*. For 10X Visium,the distance between the adjacent spots is 100 *μm*, and the diameter of each spot is 55 *μm*. For latent space visualization, we used a public dataset “NCT-CRC-HE-100K”^77^, which has patch-level labels for colorectal cancer HE slides. The TCGA-CRC and TCGA-BRCA cohort in The Cancer Genome Atlas (TCGA) program were used in the prognosis prediction, which contains tissue specimens and corresponding clinical data of CRC patients and breast cancer patients from several research centers (Supplementary Table S8 and Table S9).

### Target genes selection

The spatial expression pattern prediction of genes was formulated as a Multi-target regression (MTR) problem. The prediction target *Y* was a vector with the expression level of many genes. Considering that the total number of genes detected in each tissue varied and was high-dimensional, we selected a subset of the total detected genes as our prediction target. The selection criteria consisted of both gene’s spatial expression patterns and expression proportion across all spots. We used SPARK-X^78^ method to select spatially variable genes (SVGs) as target genes. We then choosed SVGs expressed in more than 30% spots across all tissue sections. Specifically, we performed the following steps. Firstly, we calculated SVGs for all tissues in the leave-one-patient-out validation set. For tissues from the same patient, we took the union of their SVGs. Then, we selected the intersection of the SVGs for all patients. After filtering out genes with expression proportions less than 30%, we finally selected target genes for subsequent analysis. For breast cancer, cSCC, and CRC, we included 321, 487, and 323 target genes for analysis and modeling.

### Gene expression data preprocessing

The gene expression level in spatial transcriptomics was represented as observed count data, with the key feature of over-dispersion. We transformed the raw counts data to make *Y* approximately follow a normal distribution, then mean squared error (MSE) can be used as the loss function. The first step was pseudo counts construction. We constructed the pseudo counts by adding one to each gene expression data to avoid zero values being used in the log transformation. The second step was spot-level normalization. For each spot, the pseudo counts for each gene were divided by the total pseudo counts across all genes in that spot. The third step was scaling, which multiplied all normalized count values by 1,000,000. The final step was transforming pseudo counts since the gene expression counts were highly skewed.

### HE-stained image preprocessing

We segmented each HE-stained histological image into many patches according to the coordinates of the spot. Each patch was resized into 200×200 pixels, with a 0.5 *μm*/pixel resolution. We then performed Reinhard color normalization^79^ for all HE-stained histological images to eliminate differences between stains of different tissue slides.

### Nuclei segmentation and features extraction

We used the HoVer-Net model^80^ trained on an open pan-cancer histology dataset PanNuke^81^ to segment and classify the nuclei in our HE-stained histological images of colorectal cancer. HoVer-Net can output the contour position of the nucleus and classify them into six types: background, neoplastic, inflammatory, connective, dead, and non-neoplastic epithelial. We used the Python package HistomicsTK* to perform standard color deconvolution for each color-normalized patch and then obtain their hematoxylin-stained nuclei channel. Based on the nucleus mask and the nuclei channel of patches, we can obtain a set of numerical features for each nucleus, including morphometry (size and shape), Fouried shape descriptor (FSD), intensity, gradient, and haralick features. The feature dimension is 79, and the calculated feature names and their variable types are described in Table S11. We standardized these numerical features by removing the mean and scaling to unit variance at the patient level. Moreover, we used one-hot encoding to map the predicted label of each segmented nucleus into a 6-dimensional vector. For example, neoplastic was represented as [0, 1, 0, 0, 0, 0]. As a result, 85 features were extracted for each nucleus.

### Nuclei-Graph construction

For each patch, we constructed a Nuclei-Graph *G* = (*𝒱, ℰ*) to characterize the geometric topology between cells in the tissue, with nodes *v*_*i*_, *v* _*j*_∈*𝒱* and edges (*v*_*i*_, *v* _*j*_)∈*ℰ*. Each segmented nuclei was represented as a node of the graph, with 85 extracted features in the **Nuclei segmentation and features extraction** section as node features. As shown in previous studies^82,83^, the edge (*v*_*i*_, *v* _*j*_)∈ *ℰ* was established deterministically if the distance *d*(*i, j*) between two nuclei is less than 20 *μm* by considering cell spatial composition and cell-cell communication, which is equivalent to 40 pixels for the patches in our study. In this case, each Nuclei-Graph constructed from one patch can be represented in the form of (**X, A**), where **X**∈ℝ^*N*⇥85^ is the 85-dimensional feature matrix with *N* nodes in the Nuclei-Graph, and **A**∈ℝ^*N*⇥*N*^ is the adjacency matrix of the graph.

We calculated the Euclidean distance between any pair of nuclei and then used this distance to define the edge connection in the Nuclei-Graph according to the equations (1) and (2) shown below.

For the *i*^*th*^ and *j*^*th*^ nucleus with Cartesian coordinates (*x*_*i*_, *y*_*i*_) and (*x* _*j*_, *y* _*j*_) (which use pixels as the unit) in the same HE-stained hematological image, their Euclidean distance can be calculated as follows:

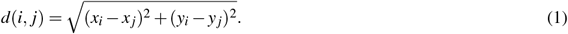

The edge connection between the *i*^*th*^ and *j*^*th*^ nucleus was assigned in the *a*_*i, j*_ element of adjacency matrix **A** as follows:

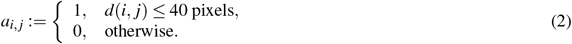

where 0 denotes that there is no edge between these two nodes, and 1 denotes that there is an edge.

### Architecture of designed IGI-DL

We proposed an integrated graph and image deep leaning model to predict gene spatial expression pattern based on HE slides. Figure 1b outlines the architecture of our designed model, which contains two input branches and one fusion output branch. The graph branch uses a four-layers Graph Isomorphism Network (GIN)^84^ to aggregate neighbor node information in Nuclei-Graph, and adopts Max & Mean global pooling to compute graph-level representation of each GIN layer. The special neural message passing operator of GIN aggregation can be represented as

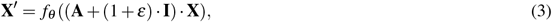

where **X** ℝ^*N*×*d*^ denotes the *d*-dimensional input node feature matrix, **A** ℝ^*N*×*N*^ is the adjacency matrix, and **I** ℝ^*N*×*N*^ is the identity matrix. *e* is a parameter, which is set as untrainable with initial value of 0 in our model. **X**^0^ ℝ^*N*×*H*^ is the aggregated *H*-dimensional output node feature matrix. *f*_Q_ denotes a neural network with parameters *q*, which is a Multilayer Perceptron (MLP) in our designed model architecture. For each node in the Nuclei-Graph, the feature aggregation process can be represented as (4), where the set of neighborhoods of a node i is denoted as *N* (*i*).

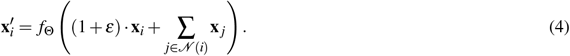

Max global pooling and mean global pooling both generate graph-level outputs by adding and averaging node features across the node dimension, respectively. Their output *r*_*Max*_ and *r*_*Mean*_ are computed by equation (5) shown below:

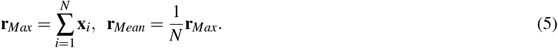

Two fully connected layers with ReLU activation^85^ are then used to integrate the graph features **r**_*Max*_⊕**r**_*Mean*_∈ℝ^2\**H*^ summed across different layers and obtain higher-level geometric information of the input Nuclei-Graph with size ℝ^*H/*2^.

The image branch uses the deep convolutional neural network that contains convolutional layers of ResNet18^61^ to extract detailed tissue texture information of HE patches with size of 200×200 pixels and applied the average pooling to output a feature vector of size ℝ^512^.

For the fusion output branch, the geometric features extracted from the graph and the texture features extracted from the image are concatenated together. The fused feature vector is then passed through with three hidden layers, and finally output predicted gene expression vector. The detailed parameters of our designed integrated model are tabulated in Table S12.

### Architecture of comparison models in the ablation experiment

In the ablation experiment for IGI-DL, we compared our designed integrated model with image-based models and graph-based models. For image-based models, there are different strategies to handle with image pixel matrices, such as convolutional operations and transformer encoders. We tested the performance of using ResNet18 and ViT as image feature extractor, respectively. Image-based ResNet18 uses convolutional layers of ResNet18 without pretraining to extract features. Image-based ViT splits each 200×200 pixels HE image into fixed-size 10×10 pixels tiles, then linearly embeds each of them, adds position embeddings, and puts each tile vector into a Transformer encoder. Then these extracted features by ResNet convolutional layers or the transformer encoder are fed into an MLP with two hidden layers to output the gene spatial expression vector finally.

For graph-based models, there are different mechanisms to deal with node feature message passing in a graph. We tested three models with varying node feature aggregation function, including the GIN used in the **Architecture of the designed integrated graph and image deep learning** section, Graph Convolutional Networks (GCN)^86^, and Graph Attention Networks (GAT)^87^. GCN aggregates node features by equations (6)), which do not have an extra MLP to generate the outputs compared with GIN. In each layer, a shared linear transformation, parametrized by **W**∈ℝ^*H*×*d*^, is applied to every node with *d*-dimensional features to output *H*-dimensional node hidden features.

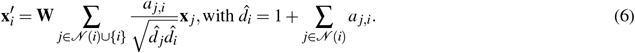

GAT uses an attention mechanism to assign neighboring nodes with different attention coefficients in the process of node features aggregation, which can be represented as

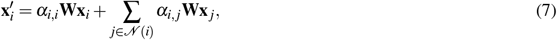

Attention coefficients *a*_*i, j*_ are computed by equations (8)). The attention mechanism is a single-layer neural network, parametrized by **a** ∈ ℝ^2\**H*^ and applying the LeakyReLU nonlinearity function (angle of the negative slope = 0.2). 3^*T*^ represents transposition and k represents the concatenation operation.

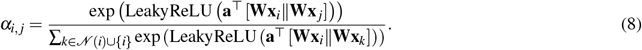

Except for neural message passing operation, the other parts of graph-based models are consistent with the graph branch in the integrated model.

For comparison, we also tested the performance of the integrated models with different combinations, including GCN+ResNet18, GAT+ResNet18, and GIN+ViT. All architecture parameters of these comparison models are listed in Table S12.

### Model training strategy

The IGI-DL and other comparison models in the ablation experiment were all trained with a mean squared error (MSE) loss function using the Adam optimizer^88^. To avoid overfitting, we performed the strategies of weight decay and early stopping. For training samples, we randomly sampled 20% into the validation set. The training process was interrupted when the loss score of the validation set stopped decreasing for specific epochs, which is set as a hyperparameter *Patience*. Detailed hyperparameter settings of different models are listed in Table S13.

Our models were implemented using PyTorch^†^ and PyG ^‡^ package (Python 3.8, PyTorch 1.10.2, PyG 2.0.3). The training and inference process were carried out on the NVIDIA HGX A100 node (40GB) of the Siyuan Mark-I cluster supported by the Center for High Performance Computing at Shanghai Jiao Tong University.

### Comparison with previous SOTA models

We compare the Pearson correlation of all genes predicted by IGI-DL with other SOTA models using the one-tailed Wilcoxon signed rank test. All SOTA models, including ST-Net, HisToGene, and Hist2ST, were trained with strategies described in their papers.

### Pathway and process enrichment analysis

We conducted pathway and process enrichment analysis based on best-predicted genes by IGI-DL using Metascape web-based platform^89^. In this platform, for each given gene list, pathway and process enrichment analysis has been carried out with the following ontology sources: KEGG Pathway, GO Biological Processes, Reactome Gene Sets, Canonical Pathways, CORUM, WikiPathways and PANTHER Pathway. For CRC, pathway and process were enriched based on accurately predicted 53 genes (Mean Pearson correlation ≥0.25 on the leave-one-patient-out validation set). For breast cancer, pathway and process were enriched based on accurately predicted 69 genes (Mean Pearson correlation ≥0.25 on the leave-one-patient-out validation set). For cSCC, pathway and process were enriched based on accurately predicted 80 genes (Mean Pearson correlation ≥0.25 on the leave-one-patient-out validation set).

### Latent space visualization

We used two different dimension reduction methods to embed the latent space in our designed integrated model into a 2D visualization space, which are t-SNE and UMAP. We implemented dimension reduction and visualization with scikit-learn, umap-learn and matplotlib in Python (Python 3.8, scikit-learn 1.0.2, umap-learn 0.5.2, matplotlib 3.3.1).

### Super-patch graph construction for HE-stained WSI

To further perform prognosis prediction on the TCGA dataset based on the spatial gene expression predicted by our IGI-DL model, we constructed a Super-patch graph from each HE-stained WSI in TCGA dataset. As shown in the steps of Algorithm 1, we first read the WSI at the resolution of 0.5 *μm*/pixel. Then, we removed the blank background regions in the image using the Otsu thresholding algorithm^90^. For the tissue region, we divided the entire image into patches of 100×100 pixels, corresponding to an actual distance of 100 *μm*, which is consistent with the input size of our constructed gene spatial expression prediction model. Furthermore, we used IGI-DL to predict the expression of genes that were validated to have a Pearson correlation greater than or equal to 0.25. We predicted 69 genes for breast cancer and 53 genes for colorectal cancer. These predicted gene expressions were used as features for the corresponding patches. To explore the advantage of using these gene expressions as prognostic features, we also extracted patch features using two CNN models, DenseNet^60^ and ResNet18, pre-trained on ImageNet for comparison. We used the second-to-last layer features of these models as the patch features, which were of dimensions 1024 and 512 for DenseNet and ResNet18, respectively. Taking into account the overall spatial structure of the WSI and aiming to minimize the redundancy of input information, we adopted the approach proposed by Lee et al. in the TEA-graph method^25^, which merges patches based on feature similarity and constructs WSI-level super-patch graphs. As shown in Supplementary Figure S18, for each patch, we calculated the cosine similarity between its features and those of neighboring patches within a range of 5×5. We recorded the number of neighboring patches with a similarity greater than 0.75. Then, we sorted the number of similar neighboring patches in descending order, selected the super-patch node one by one, and calculated the super-patch node features. Finally, based on the selected super-patches, we assigned edges to patches within the same 7×7 patches region, and calculated edge features (*ρ, θ*) based on their relative positions in polar coordinates. Please refer to Algorithm 1 for more details.

### Graph-based survival model construction

As shown in Figure 5a, we built a graph-based survival model using the super-patch graph constructed above as input. three GAT layers with two-heads attention mechanism and 50-dimensional latent features were used to extract input graph features. There were residual connections in every two consecutive GAT layers. Mean global pooling was used to aggregate the features of all nodes at the graph level. The graph-level features before and after each GAT layer were concatenated, and the dimension of these features was reduced by a MLP with two layers to 50, which is then concatenated with the encoded clinical features. In the second-to-last layer of the model, these extracted graph features are combined with encoded clinical features, and the final output is the patient-level risk score for survival prediction.

Our optimization goal is to maximize the Cox partial likelihood^91^ in our Graph-based Cox proportional hazards(CPH) model, which can be represented as

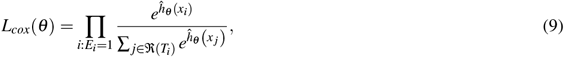

where predicted risk score *ĥ θ* (*x*_*i*_) is the output of graph-based survival model, and *θ* indicates the model’s parameters. *x*_*i*_, *x* _*j*_ are the ith and jth input super-patch graphs, respectively. *E*_*i*_ is an indicator of whether an event has occurred, with 1 representing observed death and 0 representing censored data. The risk set *j*∈𝔑(*T*_*i*_) is the set of individuals still at risk of death at time *t*.

The loss function during the training process is defined by negative log-likelihood and calculated through equation (10) shown below:

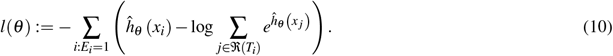

### Survival model training

In the TCGA-CRC and TCGA-BRCA cohorts, we respectively used five-fold cross-validation to train and test the aforementioned graph-based survival model, and ensured that the censorship ratio in each fold was consistent with the overall ratio. When the input data type is a spatial-gene-based super-patch graph, our graph-based survival model is lightweight as the node feature dimensions are not high, less than 100. In this case, using two 32GB V100 GPUs, the survival model can consider all samples in the training set when calculating the loss function and making optimization. On the contrary, using the DenseNet-based or ResNet-based super-patch graph as input, both the size of the input data and the parameters of the model are much larger. For TCGA-CRC and TCGA-BRCA cohorts, two and three 32GB V100 GPUs were used respectively. In order to avoid GPU memory overflowmini-batch gradient descent was used as the optimization method, with the batch size of 16 for the DenseNet-based graph and 32 for the ResNet-based graph. The Adam optimizer with learning rate of 5×10^™4^ and weight decay of 1×10^™4^ was used to optimize parameters for all these graph-based survival model. For the TCGA-CRC cohort with 559 individuals, all models were optimized 600 times, while for the TCGA-BRCA cohort with 933 individuals, all models were optimized 1000 times. Dropout rates for GAT edges, MLP layers and node/edge features were all set as 0.1 during training. In each test fold, Kaplan-Meier estimator^92^ was used to plot survival function curves. Patients in test set were divided into the high-risk group and low-risk group according to the median predicted risk score. The log-rank test was conducted to compare the difference between two survival curves of the low-risk and high-risk groups. These post-hoc statistical analyses were conducted with survival package and survminer package in R software (R 3.6.1, survival 3.1.12, survminer 0.4.9).

### IG method for survival model explanation

Integrated Gradients(IG) is an interpretability method for deep learning that can help us understand how the model makes decisions^62^. Specifically, the IG method uses the gradient information to calculate the contribution of each input feature to the model’s output. For our trained survival prediction model *ĥq* (·), the IG score of the *i*^*th*^ node feature is calculated by (11), which is integrated gradient along *i*^*th*^ dimension for an input *x* and the baseline input *x*^′^.

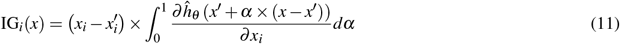

In practical computation, this integral is approximated by a summation as shown in (12), where m is the number of steps used for the summation, and the baseline input *x*^′^ is set as a super-patch graph where all node features are zero vectors. If the IG score of the *i*^*th*^ node features is positive, which means that increasing this feature will result in a higher risk score of the model output. On the contrary, negative IG score means that increasing this feature can contribute a lower risk score.

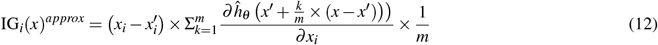

## Ethics declaration

This study was approved by the Ethics Committee of Ruijin Hospital, Shanghai Jiao Tong University School of Medicine (ID: 2020-384).

## Data availability

The spatial transcriptomics data in our in-house CRC dataset used for IGI-DL model training been deposited in the OMIX, China National Center for Bioinformation / Beijing Institute of Genomics, Chinese Academy of Sciences (https://ngdc.cncb.ac.cn/omix/: accession no. OMIX002683), which is available for non-commercial use. The spatial transcriptomics data in public CRC external test set can be obtained in Genome Sequence Archive with accession ID HRA000979. The spatial transcriptomics data in public breast cancer validation set are available at https://doi.org/10.5281/zenodo.4751624. The spatial transcriptomics data in public breast cancer test set are available at https://www.spatialresearch.org/resources-published-datasets/doi-10-1126science-aaf2403/. The spatial transcriptomics data in public cSCC validation set can be obtained in Gene Expression Omnibus (GEO, https://www.ncbi.nlm.nih.gov/geo/) with accession ID GSE144240. The spatial transcriptomics data in public cSCC test set are available at https://data.mendeley.com/datasets/2bh5fchcv6/1. The public dataset NCT-CRC-HE-100K used for latent space visualization can be downloaded from https://zenodo.org/record/1214456. The public dataset TCGA-CRC and TCGA-BRCA used for prognosis prediction can be downloaded from https://portal.gdc.cancer.gov.

## Code availability

All the related scripts and code are publicly available and can be download at https://github.com/ruitian-olivia/IGI-DL.

## Acknowledgements

We thank Prof. Pietro Liò for his helpful comments for the work. The computations in this paper were run on the Siyuan Mark-I cluster supported by the Center for High Performance Computing at Shanghai Jiao Tong University. This study was supported by National Natural Science Foundation of China (ID:12171318), Shanghai Science and Technology Development Fund (ID: 21ZR1436300), Shanghai Science and Technology Commission (20JC1410100), Shanghai Jiao Tong University STAR Grant (ID: 20190102), Medical Engineering Cross Fund of Shanghai Jiao Tong University (ID: G2021QN50).

## Author contributions

Z.Y., Y.G.W and J.S. proposed and designed the project. R.G., Y.G.W and Z.Y. developed the methodology. R.G. implemented the algorithm, ran the experiments, and wrote the first draft. X.Y., Y.M., T.W., L.J. and Y.Z. helped to analyse the data and results. Y.S., W.L., T.Z., J.Z., G.C. and J.S. acquired tissue samples and managed the spatial transcriptomics data. Z.Y., Y.G.W and J.S. supervised the project. All authors reviewed or revised the manuscript.

## Competing interests

The authors declare no competing interests.

## Footnotes

* http://github.com/DigitalSlideArchive/HistomicsTK

† https://pytorch.org/

‡ https://pytorch-geometric.readthedocs.io/

## Notes

### Competing Interest Statement

The authors have declared no competing interest.

