## supplementary materials for "Predicting Gene Spatial Expression and Cancer Prognosis: An Integrated Graph and Image Deep Learning Approach Based on HE Slides"

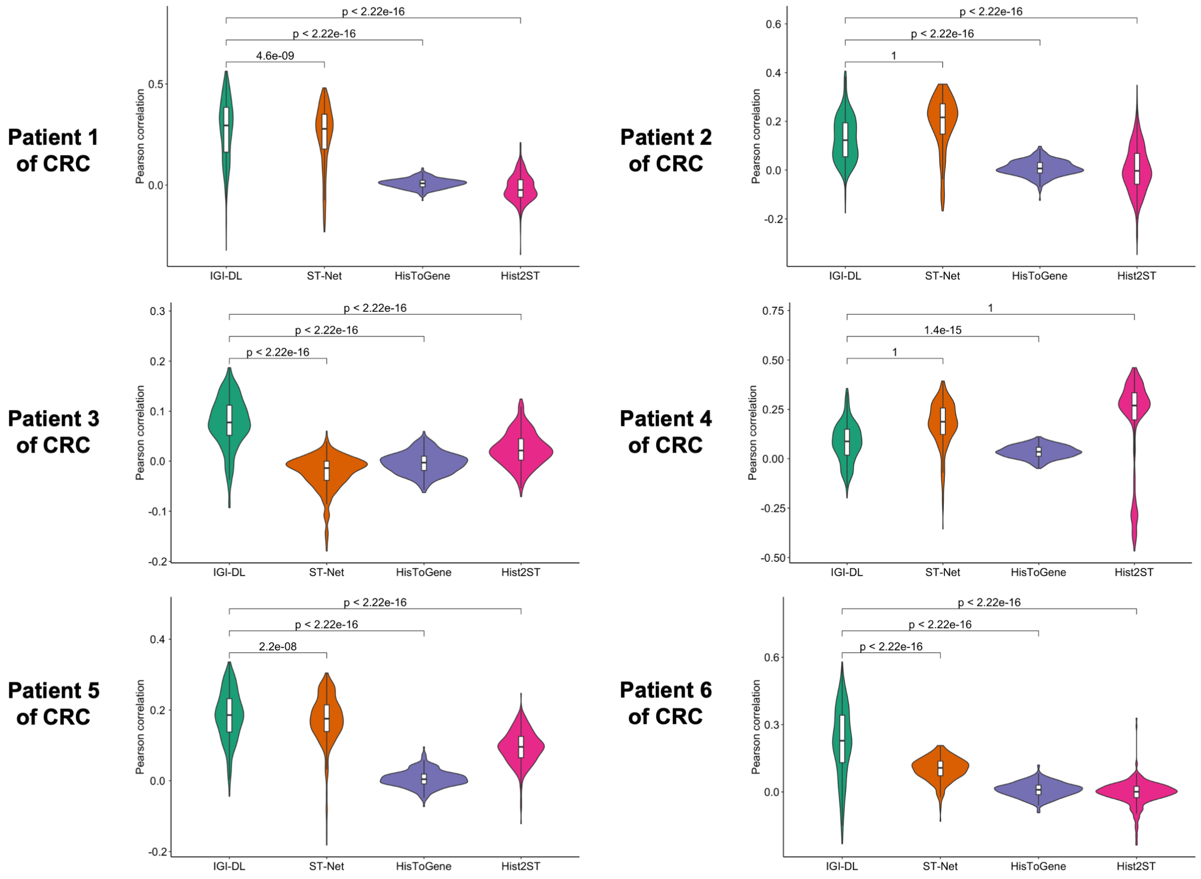


**Figure S1.** Violin plot comparing the mean Pearson correlation of target genes prediction among our model and previous models in the CRC leave-one-patient validation set.

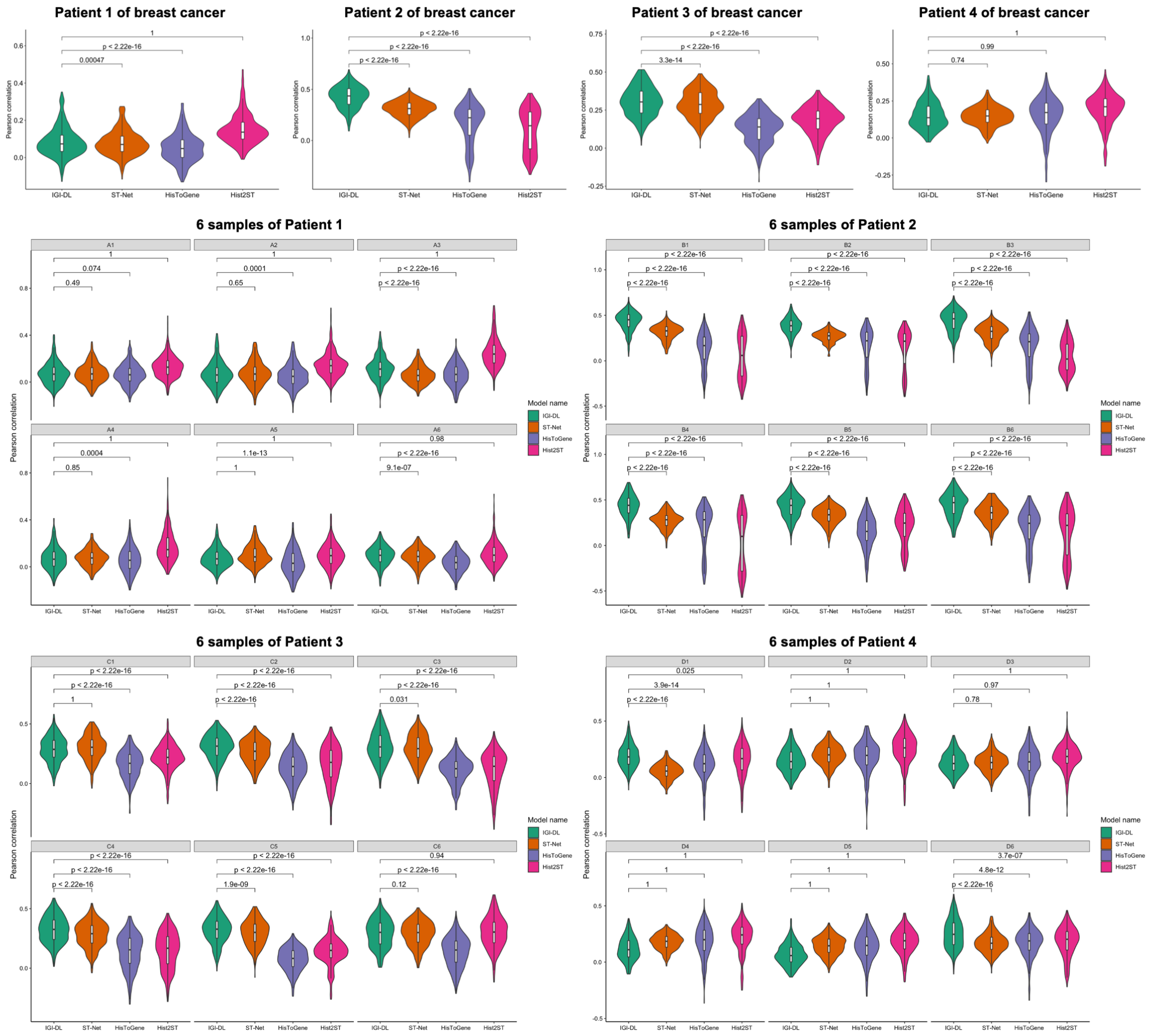


**Figure S2.** Violin plot comparing the mean Pearson correlation of target genes prediction among our model and previous models in the breast cancer leave-one-patient-out validation set (Patient 1-4).

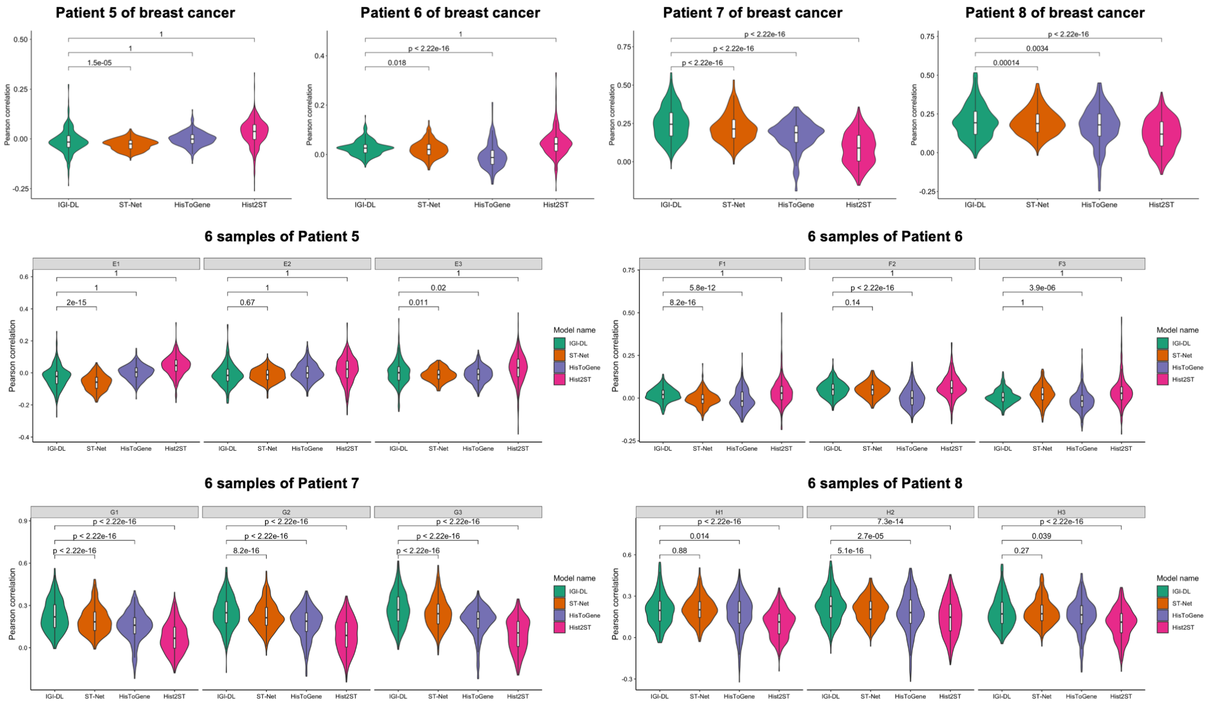


**Figure S3.** Violin plot comparing the mean Pearson correlation of target genes prediction among our model and previous models in the breast cancer leave-one-patient-out validation set (Patient 5-8).

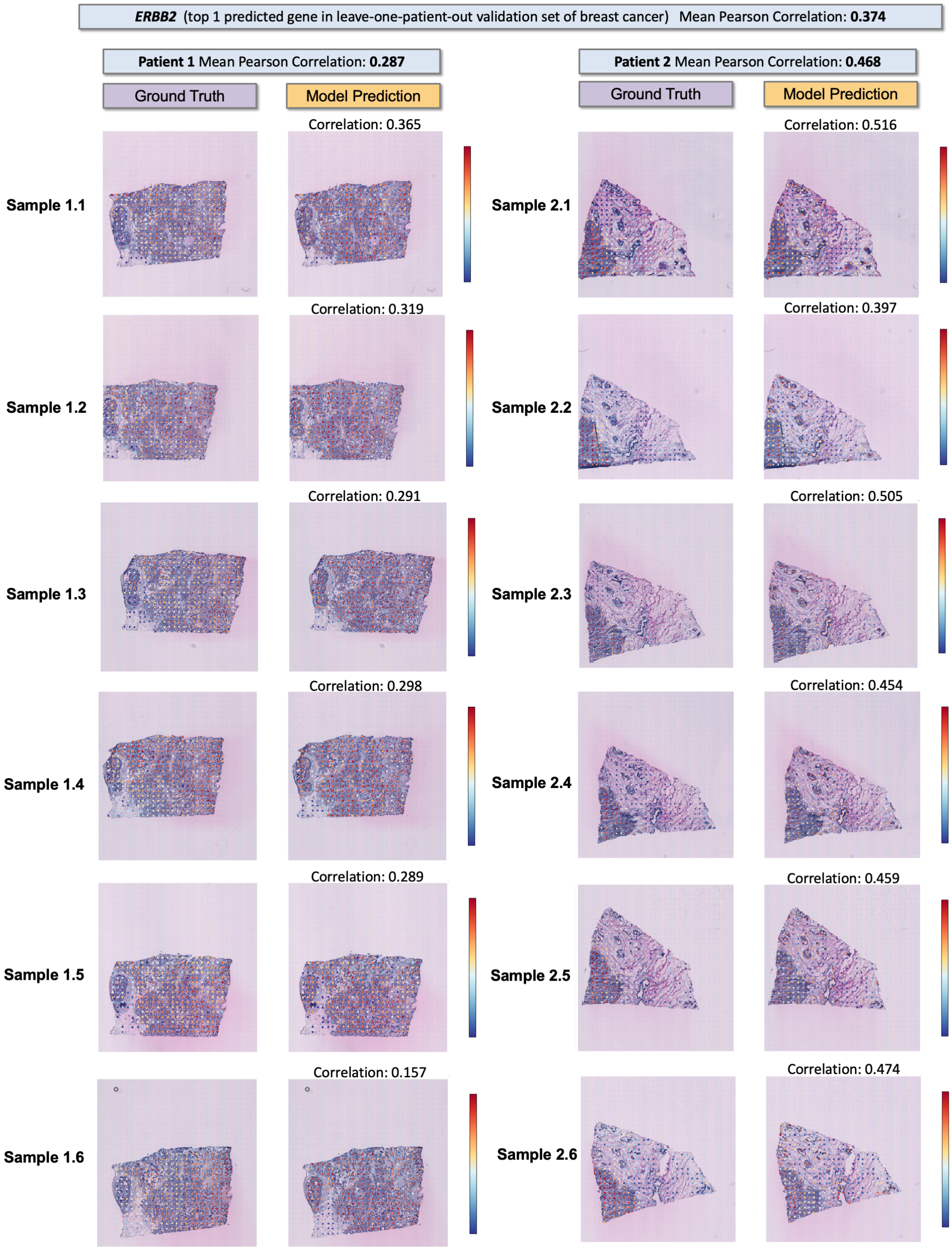


**Figure S4.** Visualization of ground-truth and predicted expression level of *ERBB2* in all tissue samples from breast cancer patient 1 and 2, where *ERBB2* ranks 1$^{st}$ among the prediction results of all target genes in breast cancer leave-one-patient-out validation set.

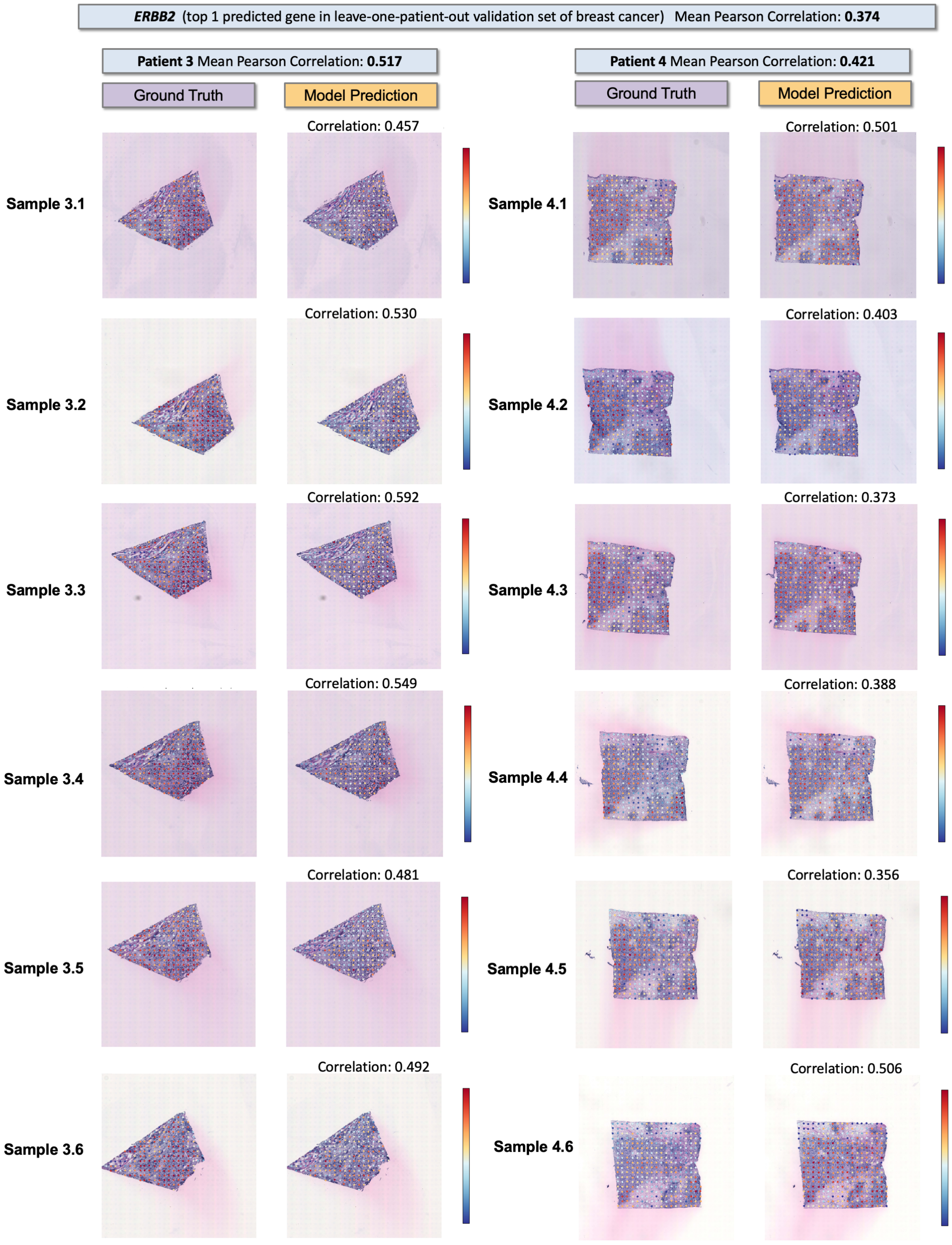


**Figure S5.** Visualization of ground-truth and predicted expression level of *ERBB2* in all tissue samples from breast cancer patient 3 and 4, where *ERBB2* ranks $1^{st}$ among the prediction results of all target genes in breast cancer leave-one-patient-out validation set.

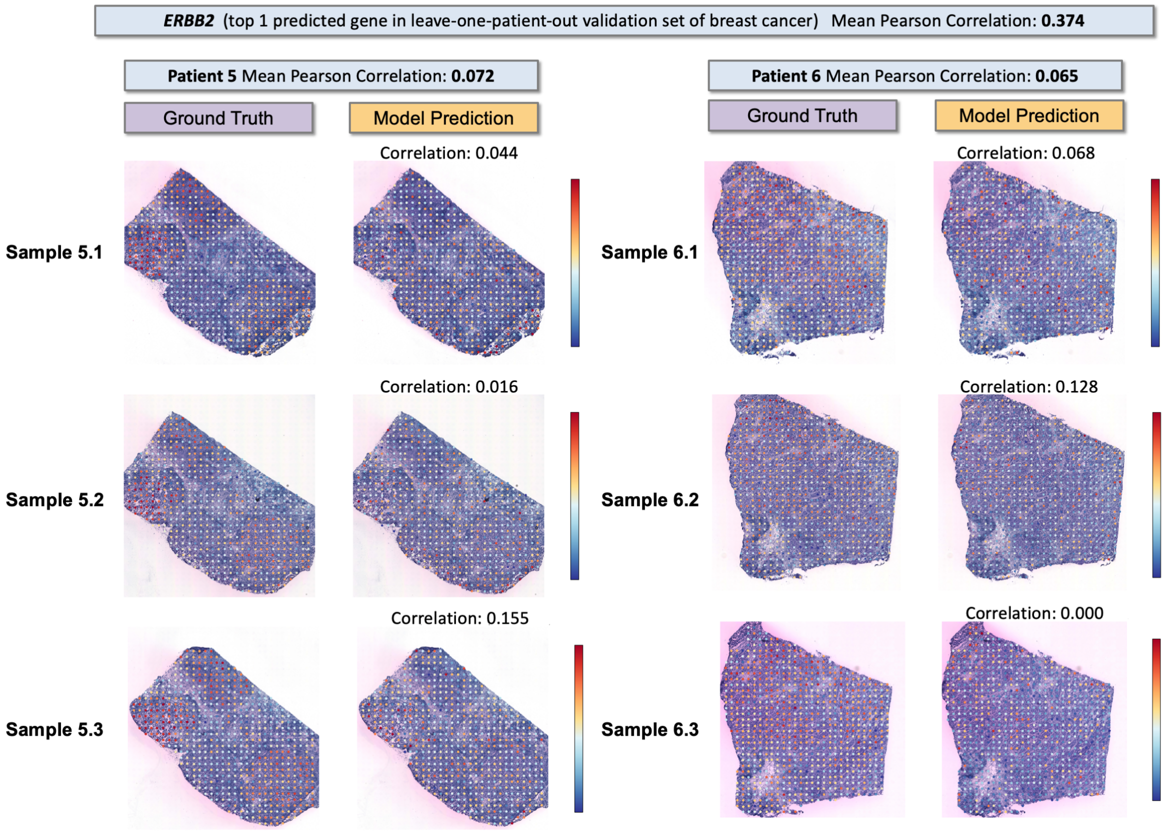


**Figure S6.** Visualization of ground-truth and predicted expression level of *ERBB2* in all tissue samples from breast cancer patient 5 and 6, where *ERBB2* ranks $1^{st}$ among the prediction results of all target genes in breast cancer leave-one-patient-out validation set.

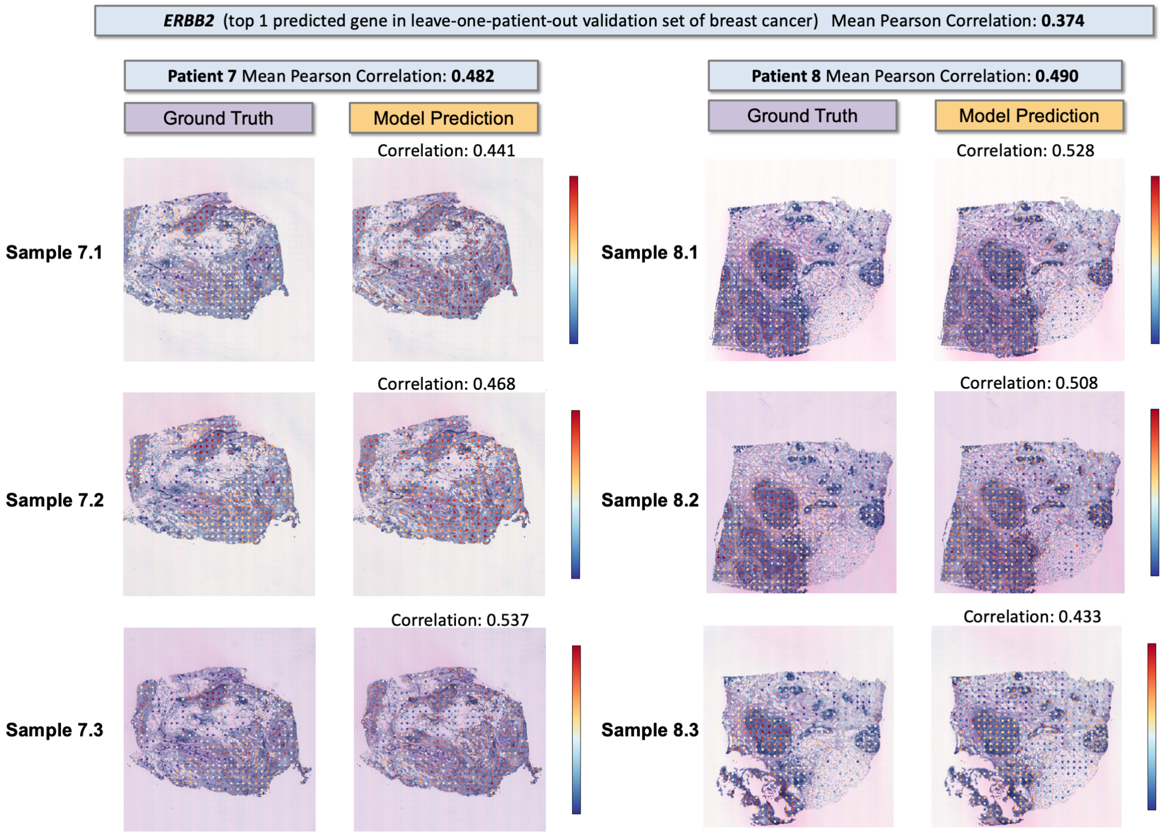


**Figure S7.** Visualization of ground-truth and predicted expression level of *ERBB2* in all tissue samples from breast cancer patient 7 and 8, where *ERBB2* ranks 1$^{st}$ among the prediction results of all target genes in breast cancer leave-one-patient-out validation set.

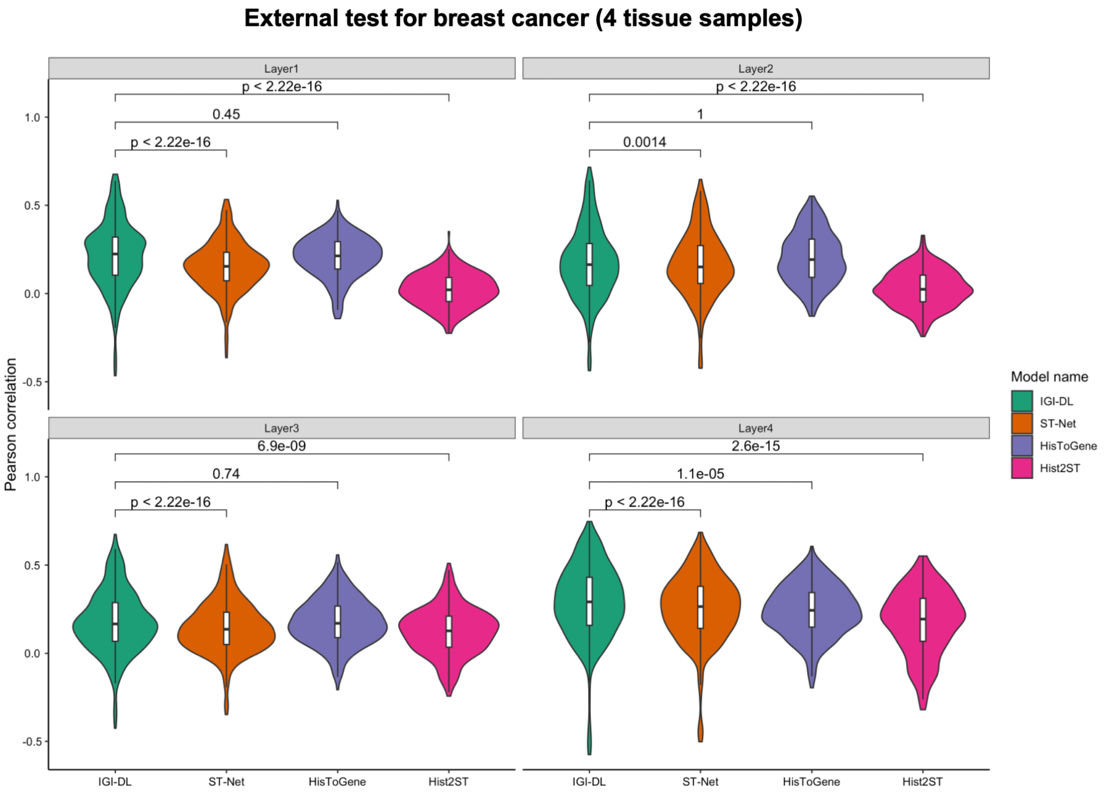


**Figure S8.** Comparison of gene prediction performance of our model and other models on each sample in the breast cancer external test set.

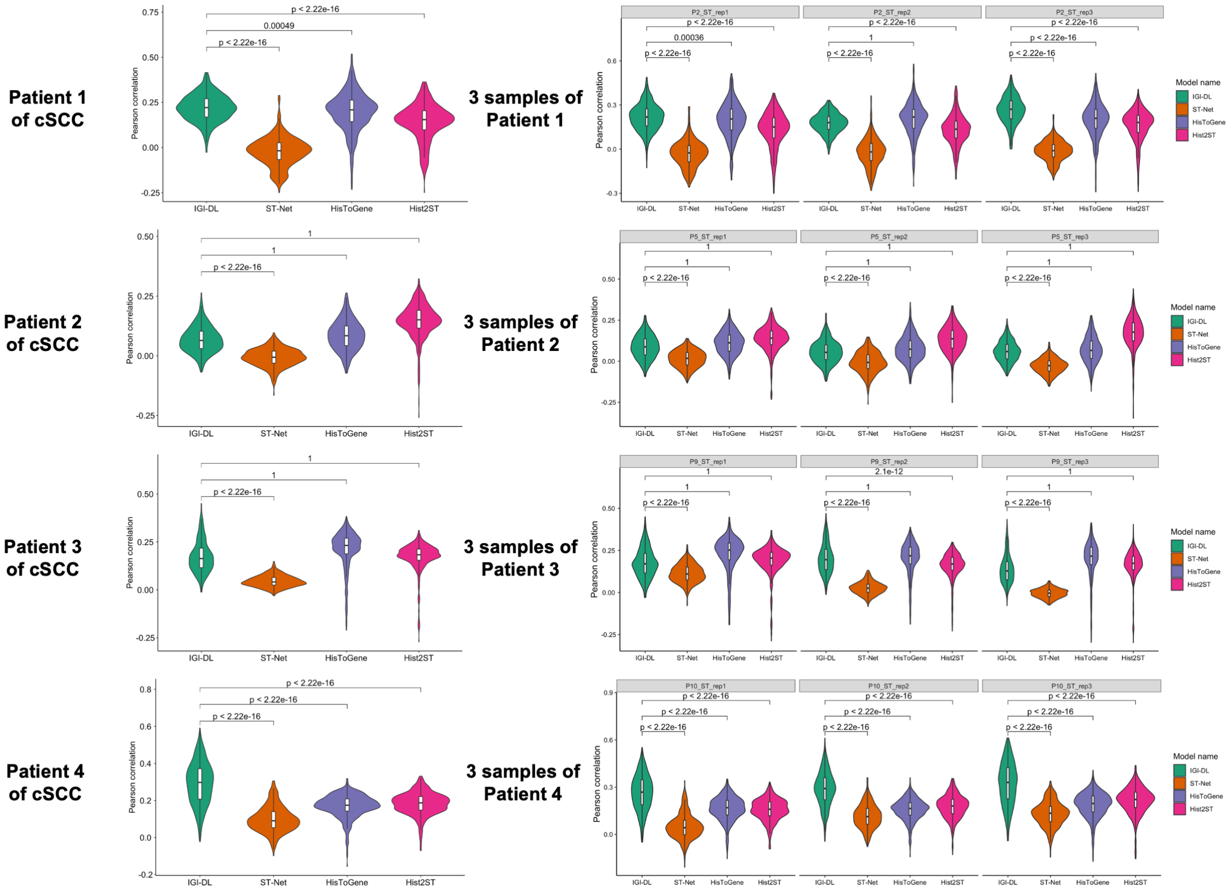


**Figure S9.** Violin plot comparing the mean Pearson correlation of target genes prediction among our model and previous models in the cSCC leave-one-patient-out validation set.

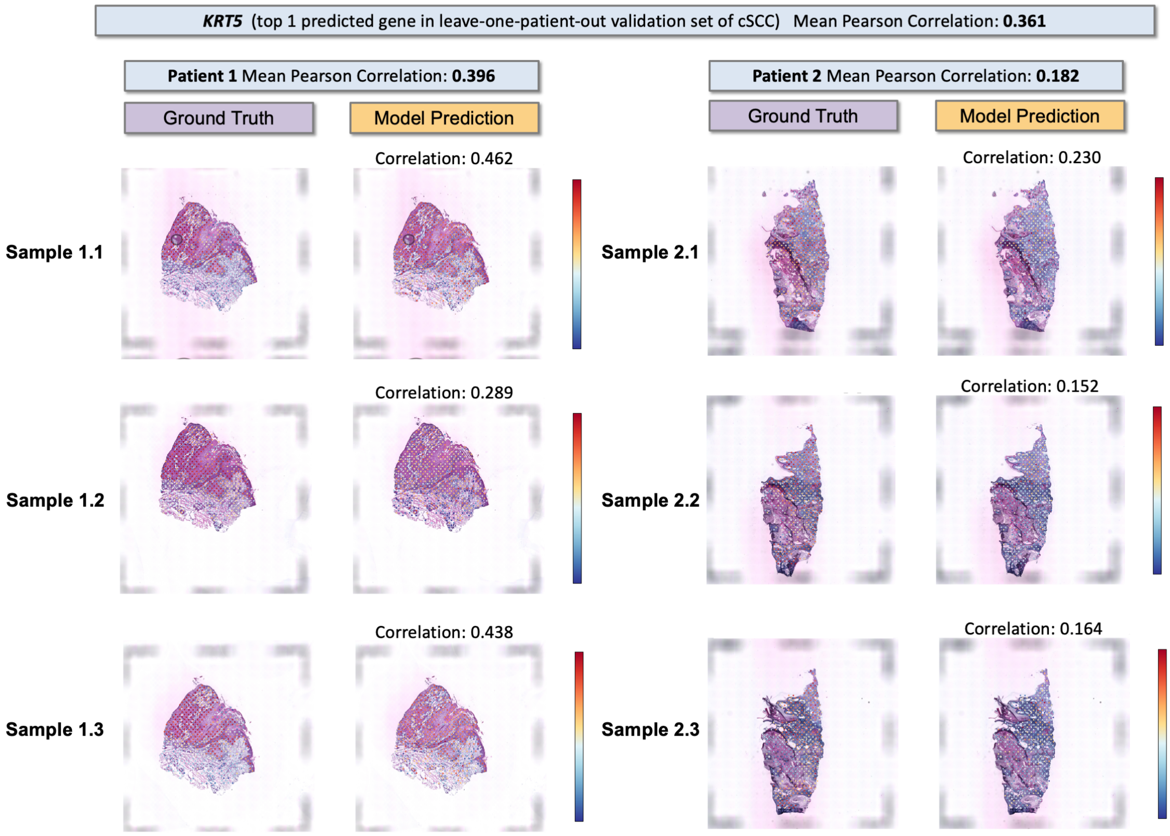


**Figure S10.** Visualization of ground-truth and predicted expression level of *KRT5* in all tissue samples from cSCC patient 1 and 2, where *KRT5* ranks $1^{st}$ among the prediction results of all target genes in cSCC leave-one-patient-out validation set.

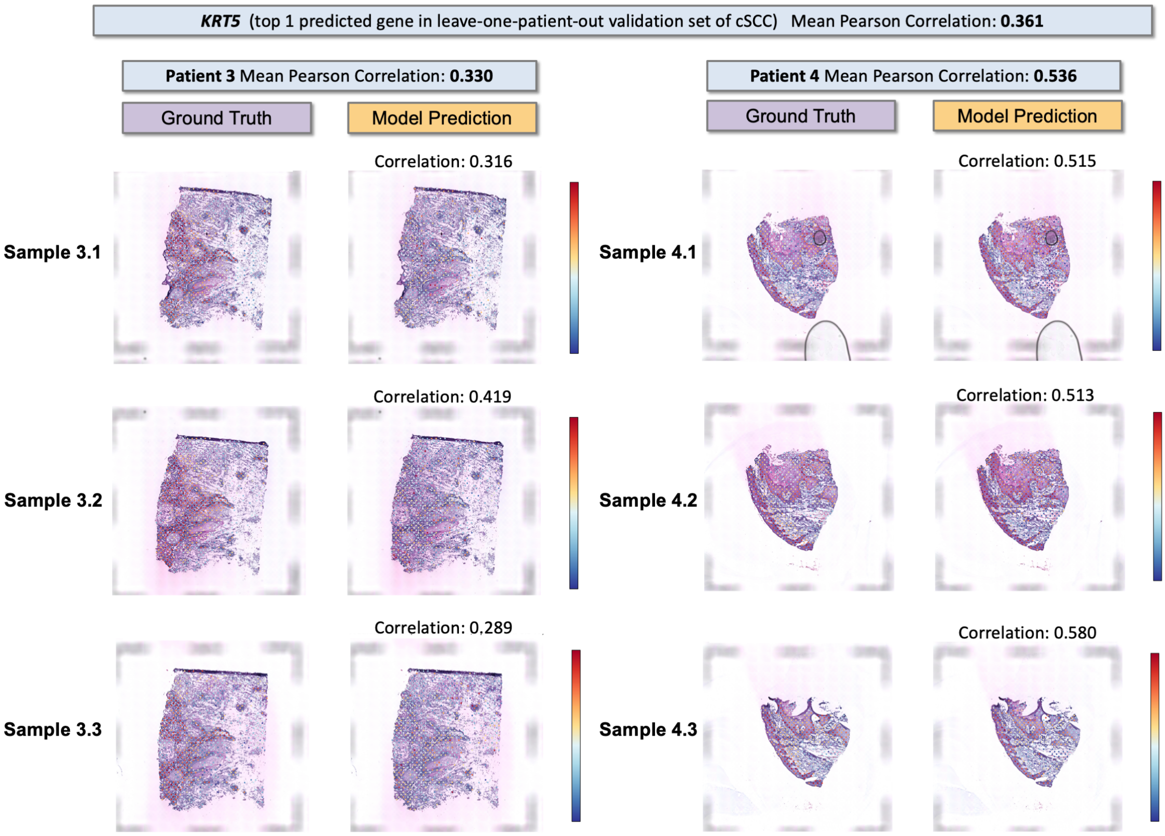


**Figure S11.** Visualization of ground-truth and predicted expression level of *KRT5* in all tissue samples from cSCC patient 3 and 4, where *KRT5* ranks $1^{st}$ among the prediction results of all target genes in cSCC leave-one-patient-out validation set.

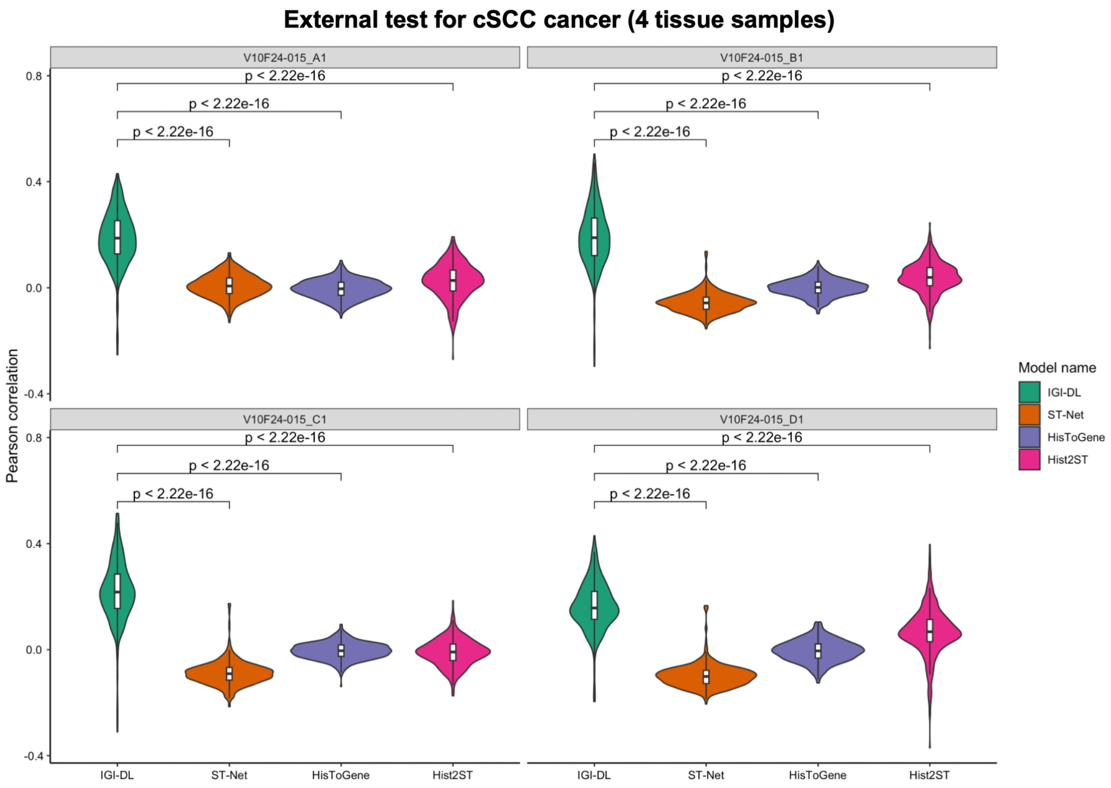


**Figure S12.** Comparison of gene prediction performance of our model and other models on each sample in the cSCC external test set.

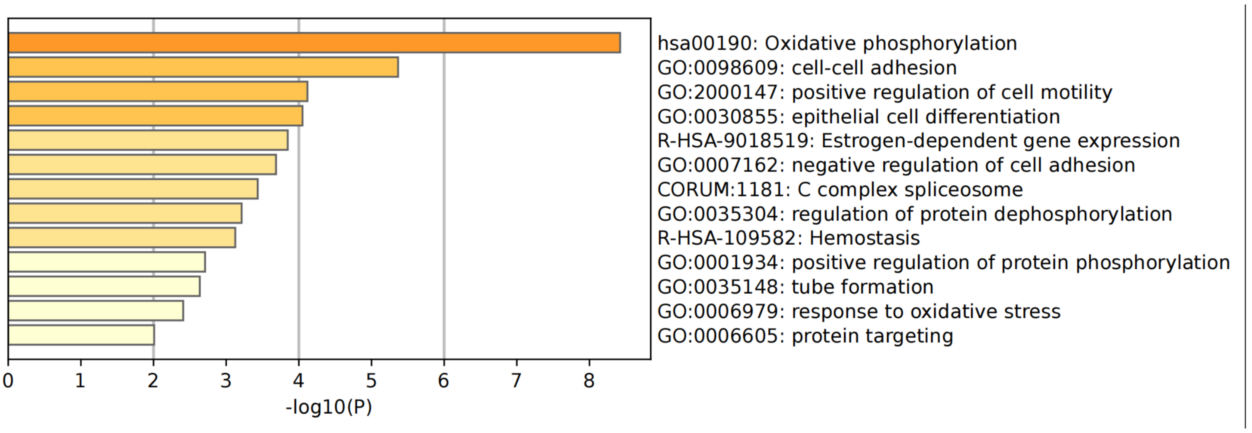


**Figure S13.** Pathway and process enrichment analysis based predicted 53 genes (mean Pearson correlation $\geq 0.25$) in CRC.

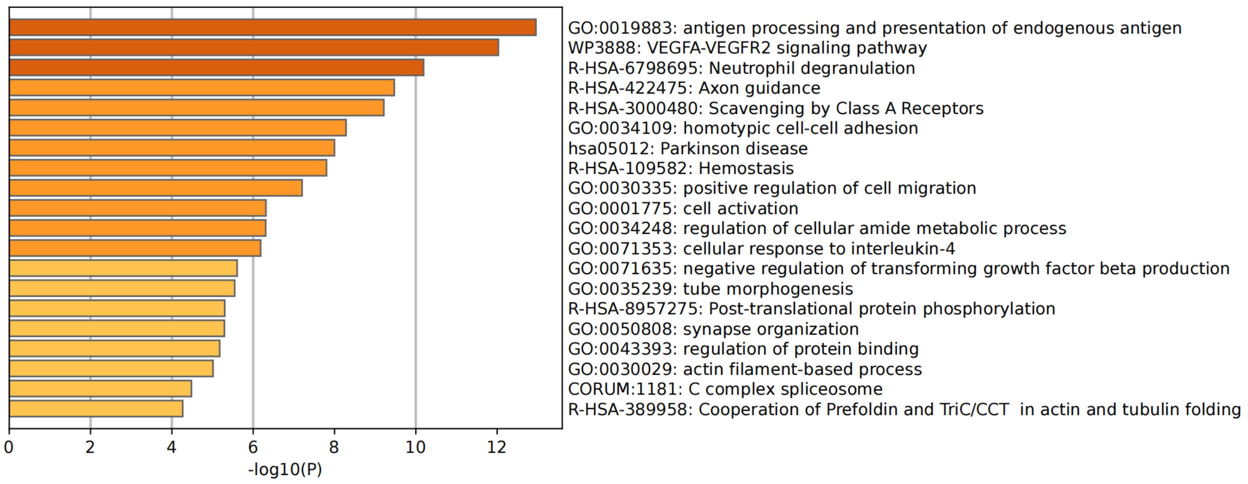


**Figure S14.** Pathway and process enrichment analysis based predicted 69 genes (mean Pearson correlation $\geq 0.25$) in breast cancer.

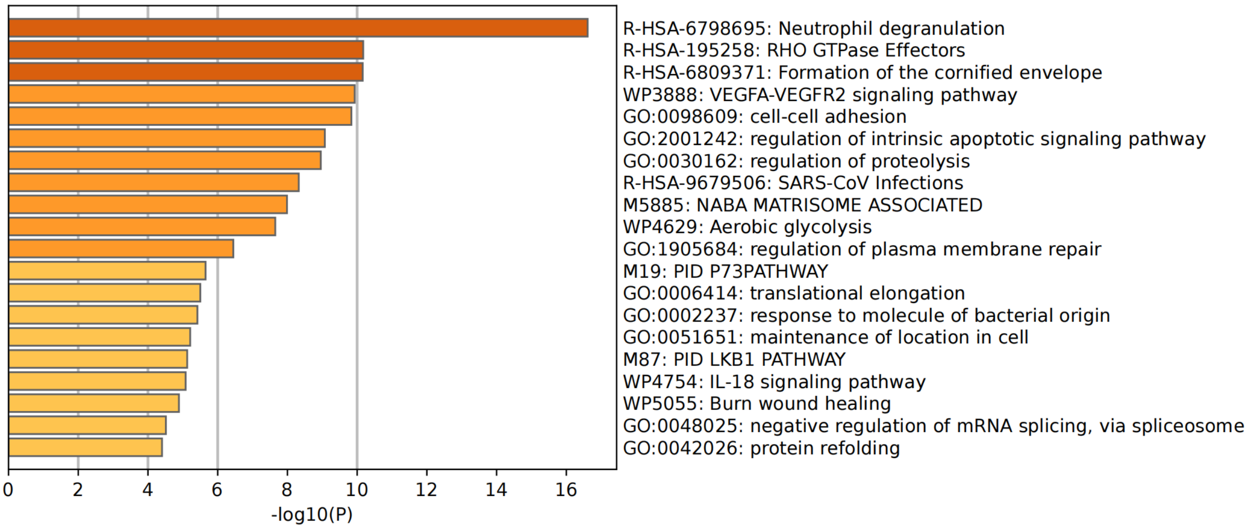


**Figure S15.** Pathway and process enrichment analysis based predicted 80 genes (mean Pearson correlation $\geq 0.25$) in cSCC.

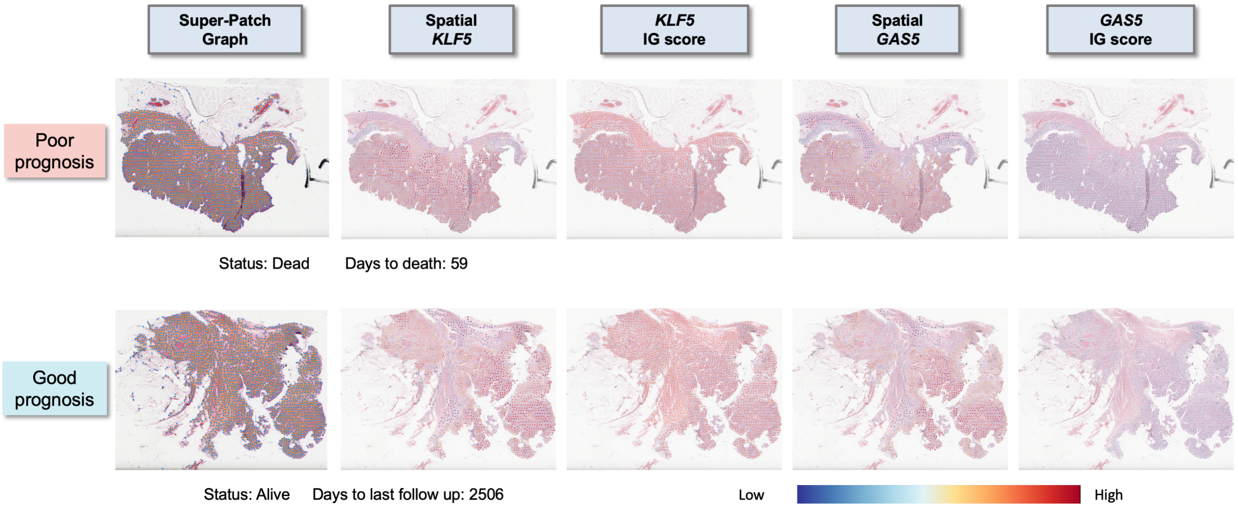


**Figure S16.** Visualization of super-patch graphs, expression patterns of unfavorable gene *KLF5* and favorable gene *GAS5* identified using the Integrated Gradients (IG) method, and their IG score of two representative tissue samples in TCGA-CRC.

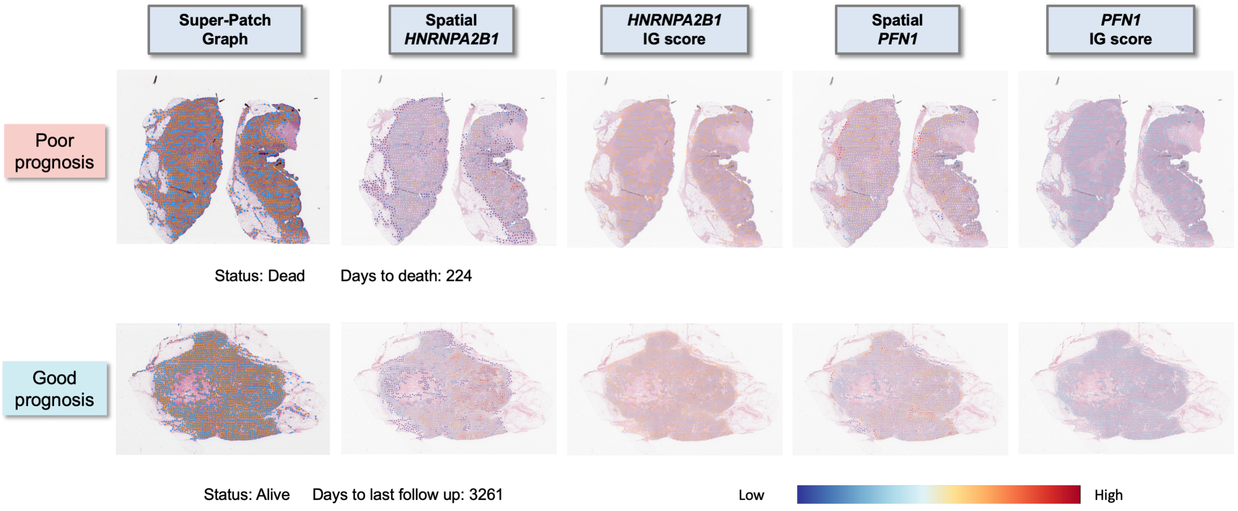


**Figure S17.** Visualization of super-patch graphs, expression patterns of unfavorable gene *HNRNPA2B1* and favorable gene *PFN1* identified using the Integrated Gradients (IG) method, and their IG score of two representative tissue samples in TCGA-BRCA.

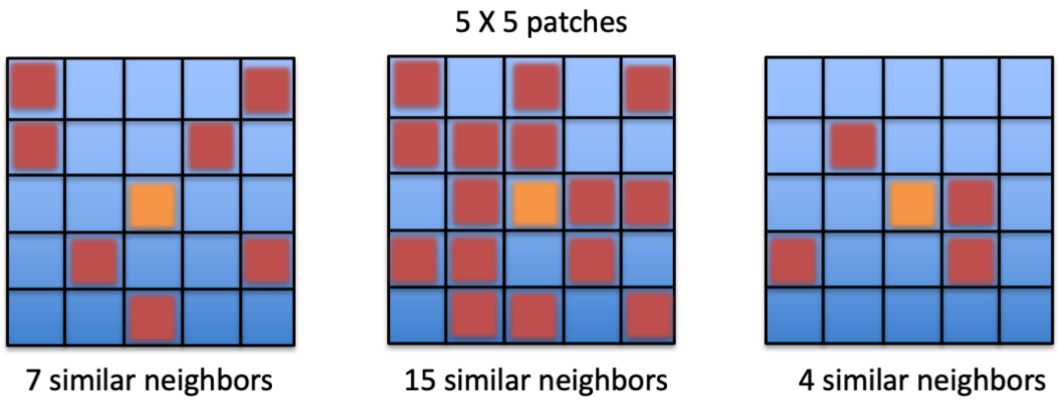


**Figure S18.** Illustration of calculating the number of neighboring patches with cosine similarity greater than 0.75 in the 5×5 patches region, where the orange patch represents the central patch and the red patches represent the similar neighboring patches that are calculated.

**Table S1.** Clinical characteristics of our in-house CRC dataset

| Patient ID | Age | Gender | Tumor Location | Stage |
|---|---|---|---|---|
| CRC1 | 32 | Male | Descending colon | IV |
| CRC2 | 58 | Male | Descending colon | II |
| CRC3 | 75 | Male | Ascending colon | IV |
| CRC4 | 54 | Male | Ileocecal valve | III |
| CRC5 | 62 | Female | Descending colon | II |
| CRC6 | 81 | Male | Rectum | II |

**Table S2.** Summary of CRC spatial transcriptomics dataset

| Cancer type | Data set | Patient ID | Sample ID | #Spots Under Tissue | #Valid Nuclei Graph |
|---|---|---|---|---|---|
| CRC | Leave-out-patient-out validation (10x Visium) | 1 | 1 | 3938 | 3909 |
| | | 2 | 2 | 4512 | 4489 |
| | | 3 | 3 | 4252 | 4195 |
| | | 4 | 4 | 3437 | 3269 |
| | | 5 | 5 | 4744 | 4714 |
| | | 6 | 6 | 3780 | 3736 |
| | External test (10x Visium) | 1 | 1 | 4248 | 4212 |

**Table S3.** Summary of breast cancer spatial transcriptomics dataset

| Cancer type | Data set | Patient ID | Sample ID | #Spots Under Tissue | #Valid Nuclei Graph |
|---|---|---|---|---|---|
| Breast Cancer | Leave-out-patient-out validation (ST) | 1 | 1.1 | 346 | 342 |
| | | | 1.2 | 325 | 319 |
| | | | 1.3 | 359 | 352 |
| | | | 1.4 | 343 | 340 |
| | | | 1.5 | 332 | 331 |
| | | | 1.6 | 360 | 349 |
| | | 2 | 2.1 | 295 | 283 |
| | | | 2.2 | 270 | 319 |
| | | | 2.3 | 359 | 352 |
| | | | 2.4 | 343 | 340 |
| | | | 2.5 | 332 | 331 |
| | | | 2.6 | 360 | 349 |
| | | 3 | 3.1 | 176 | 283 |
| | | | 3.2 | 187 | 183 |
| | | | 3.3 | 180 | 176 |
| | | | 3.4 | 184 | 179 |
| | | | 3.5 | 181 | 180 |
| | | | 3.6 | 178 | 178 |
| | | 4 | 4.1 | 306 | 304 |
| | | | 4.2 | 303 | 297 |
| | | | 4.3 | 301 | 301 |
| | | | 4.4 | 302 | 302 |
| | | | 4.5 | 306 | 303 |
| | | | 4.6 | 315 | 315 |
| | | 5 | 5.1 | 587 | 585 |
| | | | 5.2 | 572 | 570 |
| | | | 5.3 | 570 | 566 |
| | | 6 | 6.1 | 691 | 688 |
| | | | 6.2 | 695 | 689 |
| | | | 6.3 | 712 | 706 |
| | | 7 | 7.1 | 441 | 420 |
| | | | 7.2 | 467 | 438 |
| | | | 7.3 | 463 | 440 |
| | | 8 | 8.1 | 613 | 559 |
| | | | 8.2 | 603 | 544 |
| | | | 8.3 | 510 | 462 |
| | External test (ST) | 1 | 1.1 | 254 | 248 |
| | | | 1.2 | 251 | 242 |
| | | | 1.3 | 264 | 250 |
| | | | 1.4 | 262 | 255 |

**Table S4.** Summary of cSCC spatial transcriptomics dataset

| Cancer type | Data set | Patient ID | Sample ID | #Spots Under Tissue | #Valid Nuclei Graph |
|---|---|---|---|---|---|
| cSCC | Leave-out-patient-out validation (ST) | 1 | 1.1 | 666 | 558 |
| | | | 1.2 | 646 | 560 |
| | | | 1.3 | 638 | 516 |
| | | 2 | 2.1 | 590 | 444 |
| | | | 2.2 | 521 | 388 |
| | | | 2.3 | 521 | 410 |
| | | 3 | 3.1 | 1145 | 577 |
| | | | 3.2 | 1071 | 632 |
| | | | 3.3 | 1182 | 674 |
| | | 4 | 4.1 | 608 | 498 |
| | | | 4.2 | 621 | 508 |
| | | | 4.3 | 462 | 352 |
| | External test (10x Visium) | 1 | 1.1 | 2793 | 2635 |
| | | | 1.2 | 2496 | 2340 |
| | | | 1.3 | 2752 | 2672 |
| | | | 1.4 | 2363 | 2240 |

**Table S5.** Prediction performance for different cancer of our proposed IGI-DL and previous models

| Cancer type | Data set | Model name | Target gene number | Mean correlation | Median correlation | Ratio (corr≥0.25) | Ratio (corr≥0.15) |
|---|---|---|---|---|---|---|---|
| CRC | Leave-one-out validation | IGI-DL | 323 | **0.161** | **0.162** | **16.41%** | 54.49% |
| | | ST-Net | 323 | 0.143 | 0.157 | 3.41% | **55.11%** |
| | | HisToGene | 323 | 0.010 | 0.010 | 0.00% | 0.00% |
| | | Hist2ST | 323 | 0.048 | 0.057 | 0.00% | 0.00% |
| | External test | IGI-DL | 323 | **0.146** | **0.144** | **15.17%** | **47.06%** |
| | | ST-Net | 323 | 0.045 | 0.048 | 0.00% | 0.31% |
| | | HisToGene | 323 | 0.010 | 0.007 | 0.00% | 0.00% |
| | | Hist2ST | 323 | 0.079 | 0.082 | 0.31% | 10.22% |
| Breast Cancer | Leave-one-out validation | IGI-DL | 321 | **0.200** | **0.197** | **21.50%** | **78.19%** |
| | | ST-Net | 321 | 0.171 | 0.168 | 7.17% | 63.24% |
| | | HisToGene | 321 | 0.111 | 0.120 | 3.74% | 32.40% |
| | | Hist2ST | 321 | 0.128 | 0.134 | 4.98% | 43.61% |
| | External test | IGI-DL | 317 | **0.213** | 0.201 | **40.69%** | 63.72% |
| | | ST-Net | 317 | 0.179 | 0.168 | 31.55% | 57.10% |
| | | HisToGene | 317 | 0.208 | **0.203** | 35.96% | **67.51%** |
| | | Hist2ST | 317 | 0.087 | 0.086 | 6.31% | 30.28% |
| cSCC | Leave-one-out validation | IGI-DL | 487 | **0.188** | **0.183** | **16.43%** | **71.46%** |
| | | ST-Net | 487 | 0.029 | 0.027 | 0.00% | 0.00% |
| | | HisToGene | 487 | 0.166 | 0.176 | 3.70% | 68.38% |
| | | Hist2ST | 487 | 0.161 | 0.168 | 2.87% | 65.09% |
| | External test | IGI-DL | 469 | **0.191** | **0.192** | **26.01%** | **66.10%** |
| | | ST-Net | 469 | -0.059 | -0.063 | 0.00% | 0.00% |
| | | HisToGene | 469 | -0.003 | -0.003 | 0.00% | 0.00% |
| | | Hist2ST | 469 | 0.030 | 0.033 | 0.00% | 1.71% |

**Table S6.** Prediction performance for our in-house CRC dataset of different models in the ablation experiment

| Model type | | Mean correlation | Median correlation | Ratio (corr≥0.25) | Ratio (corr≥0.15) | # corr>0 (p-value<0.05) in all K-fold | corr>0 mean | Ratio (corr>0 significantly) |
|---|---|---|---|---|---|---|---|---|
| Image-based | ResNet18 | 0.144 | 0.142 | 8.36% | 45.20% | 184/323 | 0.189 | 56.97% |
| | ViT | 0.128 | 0.125 | 2.17% | 35.91% | 148/323 | 0.175 | 45.82% |
| Graph-based | GIN | 0.124 | 0.129 | 1.86% | 38.70% | **220/323** | 0.161 | **68.11%** |
| | GCN | 0.120 | 0.121 | 2.48% | 35.91% | 145/323 | 0.179 | 44.89% |
| | GAT | 0.116 | 0.118 | 0.93% | 30.65% | 155/323 | 0.163 | 47.99% |
| Integrated | GIN+ResNet18 | **0.161** | **0.162** | **16.41%** | **54.49%** | 202/323 | **0.209** | 62.54% |
| | GCN+ResNet18 | 0.145 | 0.143 | 5.88% | 47.06% | 201/323 | 0.180 | 62.23% |
| | GAT+ResNet18 | 0.150 | 0.155 | 12.69% | 52.01% | 197/323 | 0.200 | 60.99% |
| | GIN+ViT | 0.137 | 0.133 | 5.57% | 41.49% | 132/323 | 0.188 | 40.87% |

**Table S7.** Mean correlation of different models in the ablation experiment for the top five predicted genes in our in-house CRC dataset

| Model type | | Top 1 gene | Top 2 gene | Top 3 gene | Top 4 gene | Top 5 gene |
|---|---|---|---|---|---|---|
| Image-based | ResNet18 | 0.348(*EPCAM*) | 0.318(*KLF5*) | 0.299(*TSPAN8*) | 0.297(*CEACAM5*) | 0.296(*ELF3*) |
| | ViT | 0.334(*EPCAM*) | 0.300(*KLF5*) | 0.278(*TSPAN8*) | 0.276(*GPX2*) | 0.269(*CEACAM5*) |
| Graph-based | GIN | 0.308(*EPCAM*) | 0.279(*KLF5*) | 0.272(*TSPAN8*) | 0.267(*CEACAM5*) | 0.262(*GPX2*) |
| | GCN | 0.332(*EPCAM*) | 0.306(*KLF5*) | 0.287(*CEACAM5*) | 0.286(*TSPAN8*) | 0.273(*GPX2*) |
| | GAT | 0.277(*EPCAM*) | 0.262(*TSPAN8*) | 0.255(*KLF5*) | 0.238(*CEACAM5*) | 0.238(*NACA*) |
| Integrated | GIN+ResNet18 | **0.394(*EPCAM*)** | **0.349(*KLF5*)** | **0.336(*TSPAN8*)** | **0.329(*GPX2*)** | **0.328(*ELF3*)** |
| | GCN+ResNet18 | 0.328(*EPCAM*) | 0.304(*KLF5*) | 0.280(*ELF3*) | 0.274(*GPX2*) | 0.274(*CEACAM5*) |
| | GAT+ResNet18 | 0.371(*EPCAM*) | 0.334(*KLF5*) | 0.319(*ELF3*) | 0.318(*GPX2*) | 0.301(*TSPAN8*) |
| | GIN+ViT | 0.361(*EPCAM*) | 0.327(*KLF5*) | 0.303(*GPX2*) | 0.286(*CEACAM5*) | 0.285(*CD46*) |

**Table S8.** Clinical characteristics of the TCGA colorectal cancer patient cohort used for survival analysis

| Characteristics | | Summary(N=559) |
|---|---|---|
| Age at index | | 66.2±12.7 years |
| Gender | Male | 288 (51.5%) |
| | Female | 271 (48.5%) |
| Tumor type | Colon Adenocarcinoma (COAD) | 417 (74.6%) |
| | Rectum Adenocarcinoma (READ) | 142 (25.4%) |
| Status | Dead | 115 (20.6%) |
| | Alive | 444 (79.4%) |
| Stage | Stage I | 97 (17.4%) |
| | Stage II | 206 (36.9%) |
| | Stage III | 174 (31.1%) |
| | Stage IV | 82 (14.6%) |

**Table S9.** Clinical characteristics of the TCGA breast cancer patient cohort used for survival analysis

| Characteristics | | Summary(N=933) |
|---|---|---|
| Age at index | | 58.1±13.2 years |
| Status | Dead | 130 (13.9%) |
| | Alive | 803 (86.1%) |
| Stage | Stage I | 162 (17.4%) |
| | Stage II | 532 (57.0%) |
| | Stage III | 213 (22.8%) |
| | Stage IV/X | 26 (2.8%) |

**Table S10.** Prediction performance of graph-based survival models using super-patch graphs constructed by different patch feature extraction methods for colorectal cancer and breast cancer

| Cancer Type | Super-Patch Graph Type | five-Fold Cross-Validation C-index | | | | | |
|---|---|---|---|---|---|---|---|
| | | Fold 1 | Fold 2 | Fold 3 | Fold 4 | Fold 5 | Mean |
| CRC | Spatial gene expression | 0.788 | 0.628 | 0.746 | 0.713 | 0.689 | **0.713** |
| | DenseNet features | 0.720 | 0.598 | 0.617 | 0.685 | 0.625 | 0.649 |
| | ResNet features | 0.638 | 0.750 | 0.616 | 0.749 | 0.631 | 0.677 |
| Breast cancer | Spatial gene expression | 0.6654 | 0.806 | 0.730 | 0.779 | 0.727 | **0.741** |
| | DenseNet features | 0.7404 | 0.772 | 0.703 | 0.678 | 0.630 | 0.705 |
| | ResNet features | 0.727 | 0.737 | 0.645 | 0.668 | 0.618 | 0.679 |

**Table S11.** List of 79 feature names for each segmented nuclei and their corresponding variable types for graph nodes

| Feature type | Feature name | Variable type |
|---|---|---|
| Morphometry features | Orientation.Orientation | float |
| | Size.Area | int |
| | Size.ConvexHullArea | int |
| | Size.MajorAxisLength | float |
| | Size.MinorAxisLength | float |
| | Size.Perimeter | float |
| | Shape.Circularity | float |
| | Shape.Eccentricity | float |
| | Shape.EquivalentDiameter | float |
| | Shape.Extent | float |
| | Shape.FractalDimension | float |
| | Shape.MinorMajorAxisRatio | float |
| | Shape.Solidity | float |
| | Shape.HuMoments1 | float |
| | Shape.HuMoments2 | float |
| | Shape.HuMoments3 | float |
| | Shape.HuMoments4 | float |
| | Shape.HuMoments5 | float |
| | Shape.HuMoments6 | float |
| | Shape.HuMoments7 | float |
| | Shape.WeightedHuMoments1 | float |
| | Shape.WeightedHuMoments2 | float |
| | Shape.WeightedHuMoments3 | float |
| | Shape.WeightedHuMoments4 | float |
| | Shape.WeightedHuMoments5 | float |
| | Shape.WeightedHuMoments6 | float |
| | Shape.WeightedHuMoments7 | float |
| Fouried shape descriptor (FSD) | Shape.FSD1 | float |
| | Shape.FSD2 | float |
| | Shape.FSD3 | float |
| | Shape.FSD4 | float |
| | Shape.FSD5 | float |
| | Shape.FSD6 | float |
| Intensity features | Nucleus.Intensity.Min | float |
| | Nucleus.Intensity.Max | float |
| | Nucleus.Intensity.Mean | float |
| | Nucleus.Intensity.Median | float |
| | Nucleus.Intensity.MeanMedianDiff | float |
| | Nucleus.Intensity.Std | float |
| | Nucleus.Intensity.IQR | float |
| | Nucleus.Intensity.MAD | float |
| | Nucleus.Intensity.Skewness | float |
| | Nucleus.Intensity.Kurtosis | float |
| | Nucleus.Intensity.HistEnergy | float |
| | Nucleus.Intensity.HistEntropy | float |
| Gradient features | Nucleus.Gradient.Mag.Mean | float |
| | Nucleus.Gradient.Mag.Std | float |
| | Nucleus.Gradient.Mag.Skewness | float |
| | Nucleus.Gradient.Mag.Kurtosis | float |
| | Nucleus.Gradient.Mag.HistEntropy | float |

| | | |
|---|---|---|
| | Nucleus.Gradient.Canny.Sum | float |
| | Nucleus.Gradient.Canny.Mean | float |
| Haralick features | Nucleus.Haralick.ASM.Mean | float |
| | Nucleus.Haralick.ASM.Range | float |
| | Nucleus.Haralick.Contrast.Mean | float |
| | Nucleus.Haralick.Contrast.Range | float |
| | Nucleus.Haralick.Correlation.Mean | float |
| | Nucleus.Haralick.Correlation.Range | float |
| | Nucleus.Haralick.SumOfSquares.Mean | float |
| | Nucleus.Haralick.SumOfSquares.Range | float |
| | Nucleus.Haralick.IDM.Mean | float |
| | Nucleus.Haralick.IDM.Range | float |
| | Nucleus.Haralick.SumVariance.Mean | float |
| | Nucleus.Haralick.SumVariance.Range | float |
| | Nucleus.Haralick.SumEntropy.Mean | float |
| | Nucleus.Haralick.SumEntropy.Range | float |
| | Nucleus.Haralick.Entropy.Mean | float |
| | Nucleus.Haralick.Entropy.Range | float |
| | Nucleus.Haralick.DifferenceVariance.Mean | float |
| | Nucleus.Haralick.DifferenceVariance.Range | float |
| | Nucleus.Haralick.DifferenceEntropy.Mean | float |
| | Nucleus.Haralick.DifferenceEntropy.Range | float |
| | Nucleus.Haralick.IMC1.Mean | float |
| | Nucleus.Haralick.IMC1.Range | float |
| | Nucleus.Haralick.IMC2.Mean | float |
| | Nucleus.Haralick.IMC2.Range | float |

**Table S12.** Architecture of different deep learning models for gene spatial expression prediction in the ablation experiment.

| Model type | | #Neurons (MLP) | #Neurons (GNN) | #Neurons (ViT) | #Heads (Attention) | Head dim. |
|---|---|---|---|---|---|---|
| Image-based | ResNet18 | [512, 256, 256] | / | / | / | / |
| | ViT | [512, 256, 256] | / | 256 | 8 | 64 |
| Graph-based | GIN | / | 256 | / | / | / |
| | GCN | / | 256 | / | / | / |
| | GAT | / | 256 | / | / | / |
| Integrated | GIN+ResNet18 | [512, 256, 256] | 256 | / | / | / |
| | GCN+ResNet18 | [512, 256, 256] | 256 | / | / | / |
| | GAT+ResNet18 | [512, 256, 256] | 256 | / | / | / |
| | GIN+ViT | [512, 256, 256] | 256 | 256 | 8 | 64 |

**Table S13.** Hyperparameters of training different deep learning models for gene spatial expression prediction in the ablation experiment.

| Model type | | Batch size | Learning rate | Weight decay | Epochs | Patience |
|---|---|---|---|---|---|---|
| Image-based | ResNet18 | 512 | 0.0002 | 0.0001 | 300 | 30 |
| | ViT | 256 | 0.0002 | 0.0001 | 300 | 30 |
| Graph-based | GIN | 1024 | 0.0002 | 0.0001 | 300 | 30 |
| | GCN | 1024 | 0.0002 | 0.0001 | 300 | 30 |
| | GAT | 1024 | 0.0002 | 0.0001 | 300 | 30 |
| Integrated | GIN+ResNet18 | 512 | 0.0002 | 0.0001 | 300 | 30 |
| | GCN+ResNet18 | 512 | 0.0002 | 0.0001 | 300 | 30 |
| | GAT+ResNet18 | 512 | 0.0002 | 0.0001 | 300 | 30 |
| | GIN+ViT | 256 | 0.0002 | 0.0001 | 300 | 30 |

**Algorithm 1:** Algorithm of the Super-patch Graph construction

Read a svs file of one HE-stained WSI with the resolution of 0.5 $\mu m$/pixel;
Get the dimensions of the WSI: (Width, Height);
Obtain the number of rows needed to segment the WSI into patches of size (200, 200): $row\_num \leftarrow int(round(Height/200)) + 1$;
Obtain the number of columns needed to segment the WSI into patches of size (200, 200): $col\_num \leftarrow int(round(Width/200)) + 1$;
**for** $i = 1, 2, \dots col\_num$ **do**
  **for** $j = 1, 2, \dots row\_num$ **do**
    Obtain the patch for the corresponding coordinate regions;
    Calculate the proportion of the patch mask using the Otsu thresholding: $mask\_p \leftarrow thresh\_otsu(patch(i, j))$.
    **if** $mask\_p > 0.75$ **then**
      Compute the gene spatial expression for the corresponding patch using our IGI_DL model: $spatial\_gene \leftarrow IGI\_DL(patch(i, j))$;
      Save the gene expression features and record the coordinate position {X:i, Y:j} of the patch;
    **else**
      Pass;
    **end if**
  **end for**
**end for**
Calculate the number of saved patches: $patches\_num$;
**for** $k = 1, 2 \dots patches\_num$ **do**
  Initialize the number of neighboring nodes for $patch_k$: $neighbor\_patch_k = 0$;
  **for** $patch_n$ within a $5 \times 5$ patch window centered on the $patch_k$ **do**
    Measure the cosine similarity of gene expression features between two patches: $cosine\_similarity(patch_n, patch_k)$;
    **if** $cosine\_similarity(patch_n, patch_k) > 0.75$ **then**
      $neighbor\_patch_k \leftarrow neighbor\_patch_k + 1$;
    **else**
      Pass;
    **end if**
  **end for**
**end for**
Save all patches in a queue $Q$;
Sort $Q$ in descending order based on the number of neighboring nodes $neighbor\_patch_k$;
**for** $patch_k$ in $Q$ **do**
  Record $patch_k$ as a super-patch node;
  Compute super-patch node feature $spatial\_gene_k = \frac{1}{neighbor\_patch_k} \sum_{n \in N_{neighbor\_patch_k}} spatial\_gene_n$;
  Remove neighboring nodes from $Q$;
**end for**
Calculate the number of super-patches: $super\_patches\_num$;
**for** $i = 1, 2, \dots super\_patches\_num$ **do**
  **for** $j = i \dots super\_patches\_num$ **do**
    Calculate the physical distance $d_{ij}$ of node pair of $super\_patch_i$ and $super\_patch_j$;
    **if** $d_{ij} < distance\_threshold$ **then**
      Add an edge for the node pair;
      Calculate the patient-level normalized physical distance and angle as the edge features $(\rho, \theta)$;
    **else**
      Pass;
    **end if**
  **end for**
**end for**
Save the constructed Super-patch Graph.
